# Multi-layer transcriptomic characterization of age-related immune dynamics

**DOI:** 10.64898/2026.05.19.726397

**Authors:** Zhaozhao Zhao, Shuwen Zhao, Jun Jin, Ting Ni

**Affiliations:** State Key Laboratory of Genetics and Development of Complex Phenotypes, National Clinical Research Center for Aging and Medicine, Huashan Hospital, Collaborative Innovation Center of Genetics and Development, Human Phenome Institute, Center for Evolutionary Biology, Shanghai Engineering Research Center of Industrial Microorganisms, School of Life Sciences, Fudan University, Shanghai 200438, China; Multiscale Research Institute for Complex Systems, Fudan University, Shanghai 200433, China

## Abstract

Despite the pivotal role of mRNA isoform diversity in governing immune cell function, current investigations into peripheral immune aging predominantly focused on gene-level expression, obscuring deeper regulatory layers of transcriptome complexity. Here, we leveraged a 5′ scRNA-seq atlas comprising approximately 2.5 million PBMCs from 378 healthy donors. We demonstrate that immune aging is characterized by profound, non-linear transcriptional reprogramming that extends beyond gene-level shifts to include fine-tuned regulation of alternative transcription initiation and splice site selection. By quantifying the transcriptional activity of cis-regulatory elements, we resolved their contributions to age-related expression dynamics. Notably, we identified a subset of endogenous retroviruses that are reactivated in older individuals, some of which served as alternative promoters driving the production of chimeric transcripts. Furthermore, our analysis revealed *EDA* as a top-ranked gene consistently upregulated with age across multiple independent cohorts. Increasing *EDA* expression in in vitro-stimulated naïve CD4^+^ T cells from young individuals recapitulated aged phenotypes. This comprehensive resource elucidates the multi-layered transcriptomic landscape of the aging immune system and facilitates the identification of novel drivers of immune aging.

## Introduction

The human immune system undergoes continuous and dynamic remodeling throughout the lifespan. Elucidating these age-associated alterations in human peripheral blood mononuclear cells (PBMCs) is fundamental to deciphering shifting infection susceptibilities, divergent vaccine efficacies, and the pathogenesis of chronic age-related diseases^1,2^. The widespread adoption of single-cell RNA sequencing (scRNA-seq) has revolutionized our capacity to deconvolve the complexities of the immune system, enabling high-resolution interrogation of individual immune cells^3-5^. Furthermore, the emergence of large-scale datasets has facilitated the identification of rare, aging-accumulating immune subpopulations, unmasking the cellular and molecular underpinnings of altered responsiveness to both acute and chronic perturbations^6-9^. Ultimately, a deeper mechanistic understanding of these processes will yield novel therapeutic targets for precision immune modulation.

Gene expression is governed by multifaceted regulatory layers, encompassing cis-regulatory elements (CREs)-mediated long-range loops, alternative promoter selection, and alternative splicing^10,11^. These diverse modes significantly amplify the complexity of the human transcriptome and are intrinsically linked to the fine-tuning of immune regulation^12-16^. For instance, a lineage-specific alternative promoter serves as a molecular switch that selectively drives ST2 expression and IL-33 responsiveness in type 1 T cells, thereby orchestrating the expansion and clonal diversity of antiviral effector cytotoxic T cells^14^. Furthermore, transposable element (TE) exonization generates an alternative splicing isoform of the type I interferon receptor (*IFNAR2-S*); this non-canonical variant operates as a truncated decoy receptor to inhibit interferon signaling^15^. Concurrently, a subset of endogenous retroviruses (ERVs) has been co-opted as interferon-inducible enhancers, providing binding sites for STAT1 to regulate the interferon response^16^. Despite the pervasive influence of these distinct regulatory layers in shaping immune cell function, previous scRNA-seq studies investigating the age-related dynamics of PBMCs have primarily focused on gene-level profiling. Consequently, the profound diversity of transcript isoforms and their underlying regulatory networks have been largely overlooked, leaving fundamental questions unresolved regarding the temporal dynamics of isoform-level transcriptomic remodeling and potential drivers of immune aging.

In this study, we integrated a 5′ scRNA-seq atlas comprising approximately 2.5 million PBMCs from 378 healthy donors, spanning the entire human lifespan. Our analysis demonstrates the robust capacity of 5′ scRNA-seq to accurately resolve transcription start sites (TSSs) and splice junctions. We characterize a multi-layered landscape of transcriptional reprogramming during immune aging, revealing that age-associated shifts extend beyond gene-level abundance to include isoform-level regulation of alternative TSS usage and splice site selection. Notably, we uncovered non-linear trajectories in immune aging, punctuated by two global waves of transcriptomic remodeling centered at approximately ages 40 and 70. Beyond protein-coding transcripts, we quantified the transcriptional activity of cis-regulatory elements (CREs) and transposable elements (TEs), revealing their age-related dynamics. Finally, we identified *EDA* as a top one gene substantially upregulated with age across multiple independent cohorts. Through experimental validation, we demonstrated that *EDA* overexpression as a driver of functional decline of naïve CD4^+^ T cell responses in older individuals.

## Results

### A multi-layered resource of transcriptomic features across human immune cells

To characterize transcriptomic landscape of peripheral immune cells during human aging, we integrated 5′ scRNA-seq data from two independent cohorts: the WUSTL cohort^7^ (317 samples from 166 individuals aged 25-85 years old) and the SHPD cohort^8^ (61 individuals ranging from 0 to over 90 years old). In parallel, we re-analyzed scATAC-seq data from the CIMA cohort^3^ to investigate age-related epigenomic regulation. Bulk RNA-seq and ATAC-seq from isolated naïve CD4^+^ T cells was employed for orthogonal validation. We utilized CellHint^17^ to harmonize cell-type annotations across the 5′ scRNA-seq datasets, effectively aligning disparate definitions into a unified framework (Fig. 1B, S1A-B). Following integration, 22 consistently annotated cell types represented 92.3% and 96.3% of cells in the WUSTL and SHPD cohorts, respectively (Fig. S1C). The robustness of this cross-cohort alignment was further confirmed by the expression of canonical marker genes (Fig. S1D).

**Figure 1.**
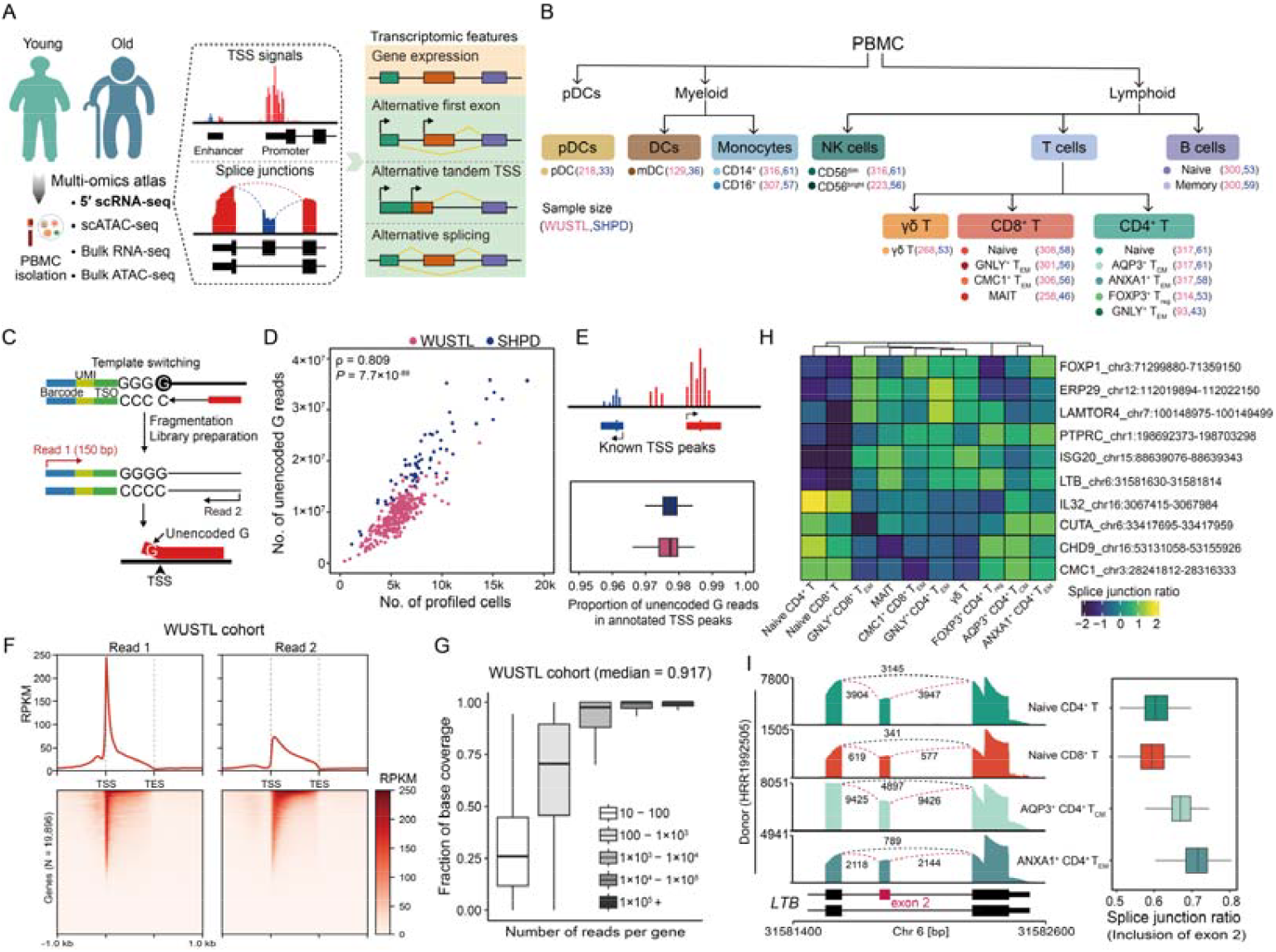
Characterizing multi-layer gene expression regulation by 5′ scRNA-seq. (A) Workflow of the overall experimental design and sample size of this study. (B) We examined 18 distinct PBMC subtypes with sufficient cell counts. Cell types are colored according to their hematopoietic lineage. The numbers located right the cell type labels indicate the sample size from two cohorts for downstream analysis. (C) Schematic overview of TSS mapping using 5′ scRNA-seq. Read 1 with the length of 150 bp sequenced beyond the template-switching oligo (TSO) and reached the actual TSS. (D) Relationship between the number of TSS reads (starting with unencoded G) and the number of profiled cells per individual across two cohorts. Pearson correlation was assessed. (E) The fraction of TSSs identified from unencoded G reads located within annotated TSS peaks per individual across two cohorts. (F) Profile plot and heatmap showing that read 1 of 5′ scRNA-seq was biased toward the TSS and read 2 was spread more evenly across the gene body. (G) The fraction of exon bases covered by reads per gene increased with the read count. The median fraction of covered exonic bases across all expressed genes was indicated. (H) Heatmap of splice junction ratios for ten representative genes undergoing alternative splicing across T cell subsets. (I) Left: Pseudobulk scRNA-seq tracks illustrating the alternative splicing of *LTB* in T cell subsets from donor HRR1992505 (SHPD cohort), with cell types and corresponding junction counts indicated. Right: Boxplot comparing the ratios of exon-included splicing junction for *LTB* across these subsets.

A pivotal advantage of the 5′ scRNA-seq data is the capacity to resolve not only gene-level expression, the primary focus of original studies, but also the dynamics of TSS selection and alternative splicing, both of which serves as fundamental mechanisms of gene regulation. By leveraging the 150-bp Read 1 sequences that span the template-switching oligo (TSO) to the cDNA 5′ termini, we precisely pinpointed TSS locations at single-nucleotide resolution^10,18^. Following the ReapTEC framework^10^, we specifically filtered for reads starting with an unencoded guanine, a hallmark of the 7-methylguanosine cap^19^ (Fig. 1C and S2A). The abundance of these TSS-containing reads (medians: 9.1M for WUSTL; 19.3M for SHPD) scaled linearly with the number of profiled cells per donor (Fig. 1D). To validate these sites, we complied 257,170 non-redundant TSS peaks from public repositories, including FANTOM5^10,20,21^. Remarkably, over 97% of our identified TSSs per donor resided within these annotated regions (Fig. 1E and S2B), underscoring the high fidelity of our detection pipeline. This sensitivity was further exemplified by the successful identification of multiple promoters for *TCF7* and *LEF1* (Fig. S2C), both of which are known to produce isoforms with different 5′ ends^22^.

We next assessed the feasibility of profiling splicing dynamics using 5′ scRNA-seq data. Consistent with previous report^4^, Read 1 exhibited a pronounced 5′ end bias, whereas Read 2 displayed a more uniform distribution across transcripts (Fig. 1F and S2D). The fractional coverage of exonic bases per gene increased with read count, reaching a median coverage of 91.7% in the WUSTL cohort and 98.1% in the SHPD cohort (Fig. 1G and S2D). Notably, reads overlapping at least one splicing junction accounted for over 50% of the total sequenced reads (Fig. S2E). Benchmarking splice junctions detected by LeafCutter^23^ against GENCODE (v44) annotations revealed that while canonical introns represented approximately 60% of unique junction sites (Fig. S2F), they accounted for over 95% of total junction-spanning reads (Fig. S2G). The reliability was further supported by recapturing lineage-specific isoform patterns in known genes. For example, the CD45RA isoform (included one or more of exons 4,5 and 6) was enriched in naïve T cells, whereas the CD45RO variant (skipping exons 4-6) predominated in activated and memory populations^13^ (Fig. S2H). Collectively, these results establish our 5′ scRNA-seq data as a robust resource for the systematic profiling of alternative TSSs and alternative splicing.

To systematically profile multi-layer transcriptomic features, we generated pseudobulk measures by aggregating TSS and splice junction counts from all cells belonging to the same cell type per donor, retaining 18 cell types with sufficient sample depth (Fig. 1B). Each pseudobulk captured a median of 8,400 expressed genes (range of medians: 6,177-9,679; Fig. S3A). Beyond gene-level expression, we quantified widespread alternative transcript isoform usage, identifying a median of 2,904 genes with multiple tandem TSSs (range: 1,359-4,353) and 414 genes displaying multiple first exons (FEs, range: 180-693) per cell type. Furthermore, we resolved a median of 3,124 alternative splicing events per pseudobulk sample (range: 996-10,558; Fig. S3A).

Leveraging this rich resource, we identified widespread differential isoform usage across T cell subtypes beyond the well-documented case of *PTPRC* (Fig. 1H). A representative example is *LTB*, which encodes a type II membrane protein of the TNF family involved in the inflammatory response and lymphoid tissue development^24^. LTB showed preferential inclusion of exon 2 in memory T cells compared to naïve T cells (Fig. 1I). Notably, exon 2 skipping introduces a frameshift resulting in a truncated protein that lacks most of the extracellular domain. Furthermore, we detected lineage-specific alternative FE usage, as exemplified by *THEMIS*, a known signaling hub critical for T cell maturation^25^, which favored an upstream FE specifically in naïve T cells (Fig. S3B and S3C).

### The landscape of aging-related transcriptomic features across immune cells

Our analytical framework provides a unique opportunity to investigate the dynamic landscape of multi-layer transcriptomic features during human aging. We first identified molecular features exhibiting age-related linear trajectories by performing Spearman correlation analysis, adjusting for potential confounding factors including Body Mass Index (BMI) and sex. To account for differences in genetic ancestry and age distribution^26,27^, the WUSTL and SHPD cohorts were analyzed independently. Across the four regulatory layers, we detected substantial genes showing age-related linear shifts, with naïve CD4^+^ T cells harboring the greatest abundance of such features (Fig. 2A). In the WUSTL cohort, for example, we identified 473 genes with age-linear expression changes, alongside 175 genes with linear alterations in tandem TSS usage, 96 genes in major FE usage, and 97 genes in splice junction ratio in naïve CD4^+^ T cells (Fig. S4A-B). Our integrated analysis expanded the catalog of aging-related signatures: the total counts of linearly changing features increased from 1,194 (expression only) to 1,839 in the WUSTL cohort, and from 1,875 to 3,601 in the SHPD cohort (Fig. 2B). We observed a positive correlation between the number of genes exhibiting age-related linear shifts in overall expression and those undergoing isoform-level changes (Fig. 2C), indicating a coordinated multi-layer program of gene regulation during aging. While the majority of age-related tandem TSS shifts primarily remodeled 5′ UTR length, approximately 50% of alternative first exon usage directly altered coding sequence (Fig. S4C). These findings suggest that dysregulated transcription initiation orchestrates the production of alternative isoforms characterized by distinct sequences, which may influence translational efficiency and protein structure in a context-specific manner.

**Figure 2.**
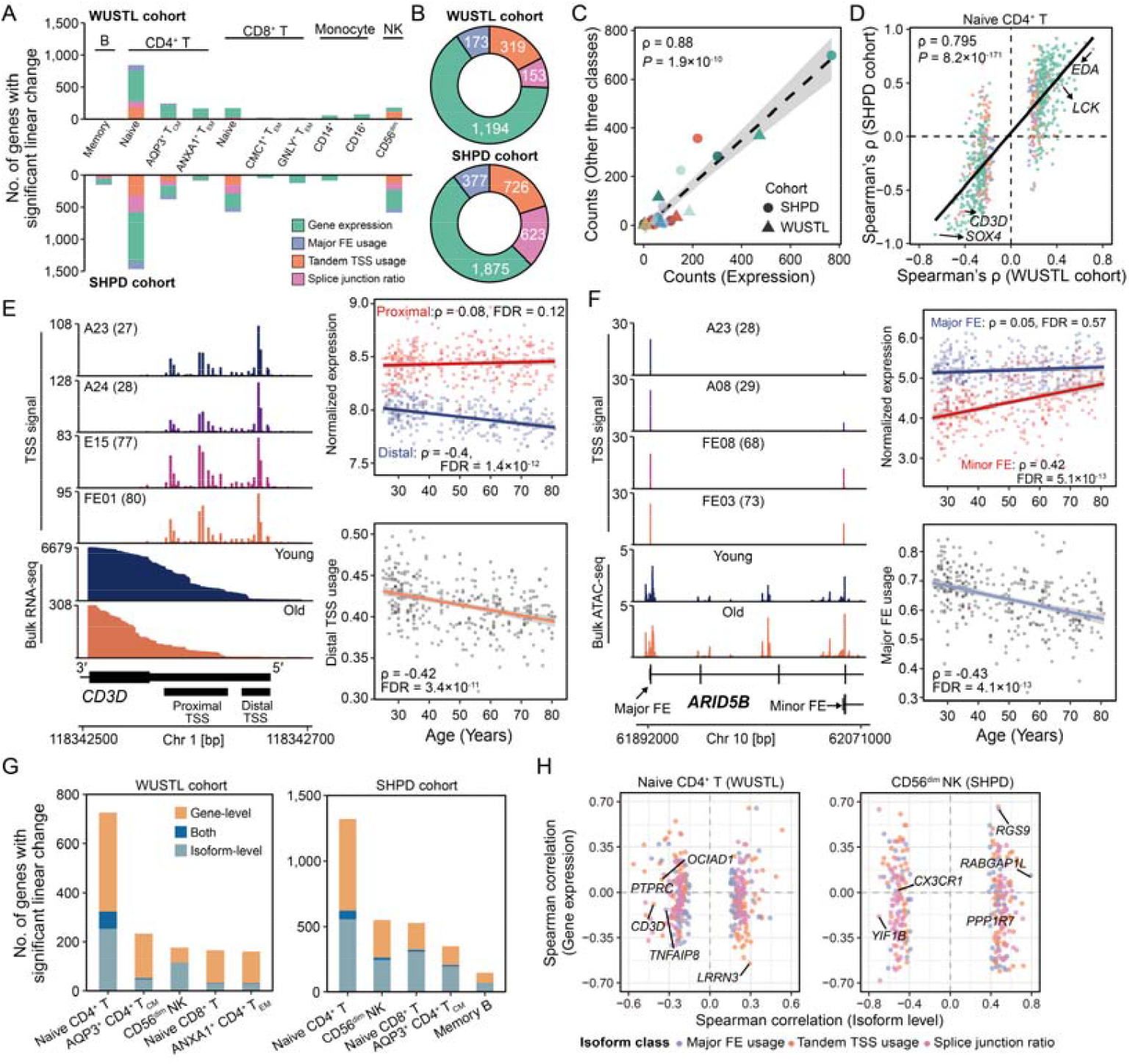
Linearly changing multi-layer transcriptomic features across immune cells during aging. (A) Number of genes with significant linear age-dynamics, stratified by transcriptomic feature layer. The ten cell subtypes with the highest abundance of linear changing features are shown. (B) Pie charts summarizing the counts of transcriptomic features showing linear changes during aging in two cohorts. (C) Correlation between the quantify of linearly changing features at the gene expression level and those at three other transcriptomic layers. (D) Scatter plot displaying the correlation of age-associated linearly changing features (stratified by transcriptomic layer) in CD4+ Naïve T cells across the two cohorts. (E) Left: TSS density profiles showing dynamic tandem TSS usage of *CD3D* during aging in CD4+ Naïve T cells. Right: Scatter plots depicting the Spearman correlation between TSS expression and distal TSS usage of *CD3D* with chronological age in CD4+ Naïve T cells from the WUSTL cohort. Each point represents an individual. (F) Left: TSS density profiles showing alternative FE usage of *ARID5B* during aging in CD4+ Naïve T cells. Right: Scatter plots depicting the Spearman correlation between FE expression and major FE usage of *ARID5B* with chronological age in CD4+ Naïve T cells from the WUSTL cohort. (G) Stacked bar chart showing the number of genes exhibiting linear changes in overall expression (gene-level), differential isoform usage (isoform-level), or both, across cell types with the most abundant transcript alterations. (H) Scatter plots showing the relationship between isoform-level Spearman correlation (x-axis) and gene-level Spearman correlation (y-axis) in CD4+ Naïve T cells from the WUSTL cohort (left) and CD56^dim^ NK cells from the SHPD cohort (right).

To evaluate the robustness of these age-associated signatures, we assess their consistency between the two independent cohorts. By correlating the Spearman’s rank coefficients derived from each cohort across all features, we observed a positive concordance across cell types, particularly within the naïve CD4^+^ T cell subset (Fig. 2D and S4D-E). For instance, we observed two tandem TSS peaks within the 5′ UTR of *CD3D* (Fig. 2E, left), a critical component of the T cell receptor complex^28^. While the proximal tandem TSS remained stable with age, the distal TSS exhibited a progressive decline in expression, leading to a significantly decreased distal TSS usage across both cohorts (Fig. 2E, right, and S4F). Similarly, *ARID5B* utilizes alternative first exons to yield two functionally distinct isoforms that differentially modulate IL-6 production^29^. While the upstream major FE maintained a steady state, the downstream FE generating a truncated isoform showed marked upregulation during aging in naïve CD4^+^ T cells (Fig. 2F and S4F). Crucially, ATAC-seq analysis corroborated this AFE dynamic, revealing increased chromatin accessibility specifically at the downstream FE of *ARID5B* with aging (Fig. S4G). Furthermore, we observed an age-related increase in the CD45RO-specific splice junction ratio of *PTPRC* within the naïve CD4^+^ T cell population, a trend that was highly consistent across both cohorts (Fig. S4B and S4F).

Notably, across both cohorts, the majority of genes exhibiting age-associated linear isoform-level shifts did not show concomitant linear changes in overall expression across cell types (Fig. 2G and 2H). In naïve CD4^+^ T cells, for example, the overall expression of *CD3D* and *PTPRC* remained stable with age in the WUSTL cohort (Fig. S5A). Similarly, in CD56^dim^ natural killer (NK) cells, the chemokine receptor *CX3CR1* exhibited a progressive age-related shift toward upstream FE usage in the SHPD cohort, despite maintaining constant total mRNA expression (Fig. S5B and S5C). Collectively, these findings demonstrate that dissecting transcript isoform dynamics uncovers a critical layer of transcriptomic regulation and novel molecular biomarkers during the aging of peripheral immune cells that were invisible to conventional gene-level analysis.

### Nonlinear trajectories of immune aging revealed by temporal clustering

To delineate the temporal dynamics of immune aging, we categorized the WUSTL cohort into discrete age groups and investigated transcriptomic dysregulation relative to a young baseline (ages 25-35 year). This analysis identified 6,380 features significantly altered in at least one age group across multiple cell types (Fig. S6A). Notably, only 22.7% (N = 1,450) of these dysregulated features exhibited a linear correlation with age, suggesting that the majority of molecular shifts follow complex, nonlinear trajectories. In naïve CD4^+^ T cells, for example, the older group (ages 65-81 year) harbored the highest burden of dysregulation across tandem TSS usage, major FE usage and overall gene expression (Fig. S6B and S6C). Conversely, the most pronounced shifts in splice junction ratios were observed in the middle-aged group (ages 45-54 year, Fig. S6B). Trajectory pattern of all dysregulated features in naïve CD4^+^ T cells further underscored widespread nonlinear temporal patterns (Fig. S6D).

To uncover specific patterns of transcriptomic features during aging, we employed unsupervised fuzzy c-means clustering to group features with similar trajectories^30^. We integrated all four transcriptomic layers across 18 cell types, yielding a total of 287,070 features. Ten distinct clusters of molecular trajectories were identified, with the number of features per cluster (membership above 0.7) ranging from 14,602 to 20,280 (Fig. 3A, 3B and S6E-S6H). Each cluster comprised features from all four regulatory layers and multiple cell lineages, reflecting a coordinated and shared transcriptomic architecture of aging across the immune system (Fig. 3A and 3B). Most trajectory patterns changed nonlinearly with age, further confirming that immune aging is a nonlinear decline (Fig. 3B).

**Figure 3.**
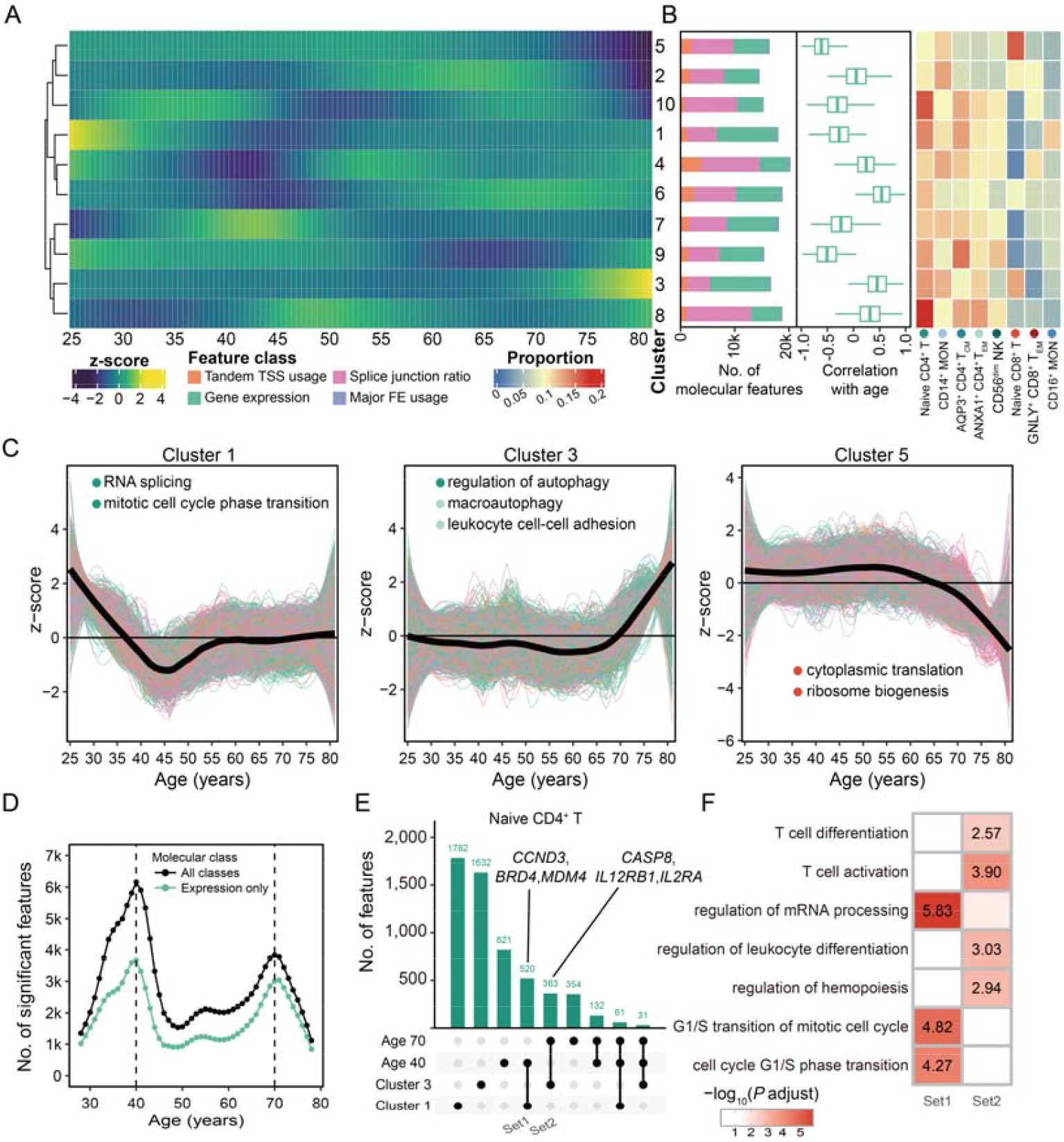
Nonlinear transcriptomic changes during immune cell aging. (A) Heatmap showing the age-associated molecular trajectories across 10 consensus clusters. All four classes of molecular feature from 18 immune cell subtypes were grouped for clustering. (B) Left: Stacked bar plot showing the compositional distribution of molecular feature classes. Middle: Box plots showing the distribution of Spearman correlation coefficients between molecular features and age. Right: Heatmap showing the proportion of molecular features coming from different cell subtypes in 10 clusters. (C) Three representative clusters of molecular features exhibiting clear and straightforward nonlinear changes during human aging. Significantly enriched GO terms were displayed, with cell subtypes indicated by colored points. (D) Number of molecular features exhibiting significant age-associated changes, with two prominent peaks observed at ages 40 and 70, depicting trends for the total number of all four feature classes and the subset of differentially expressed genes. (E) The overlapping of transcriptomic features between two ages (40 and 70) and two clusters (cluster 1 and 3) in CD4+ naïve T cells. (F) Heatmap showing the GO enrichment for molecular features within two overlapping sets.

Among these, three (Clusters 1, 3 and 5) exhibited clear and easily understandable dynamic patterns across the human lifespan (Fig. 3C and S7A). Cluster 1 decreased linearly between ages 25 and 45, then rose and stabilized after age 55; features derived from naïve CD4^+^ T cells in this cluster were enriched for RNA splicing and cell cycle pathways (Fig. 3C, left and S7). Cluster 3 remained relatively stable until age 65, followed by a sharp terminal increase (Fig. 3C, middle). In contrast, Cluster 5 maintained a plateau until approximately age 55 and then declined rapidly; naïve CD8^+^ T cell features within this cluster were functionally linked to translation and ribosome biogenesis (Fig. 3C, right).

### Uncovering waves of aging-related transcriptomic changes across lifespan

To quantitatively delineate transcriptomic dynamics across the human lifespan, we applied the DE-SWAN algorithm^31^. This approach identifies dysregulated features by progressively scanning 30-year windows and comparing molecular levels between adjacent 15-year intervals. By aggregating significantly changed features across all 18 cell types, we uncovered two prominent, global waves of transcriptomic dysregulation, centered at approximately ages 40 and 70 (Fig. 3D and S7C). These two waves aligned closely with our trajectory clustering result (Fig. 3A), suggesting a high degree of concordance between independent analytical methods. The first wave (age ∼40 year) corroborated with two recent reports, one based on scRNA-seq^9^ and the other on bulk RNA-seq^32^. This bimodal pattern remained robust across varying sliding window parameters and was consistently observed when considering only overall gene expression (Fig. 3D and Fig. S7D). Importantly, similar waves were evident across diverse immune cell types (Fig. S7E), indicating that these aging-related shifts represent coordinated and systemic alterations rather than changes confined to specific transcriptomic layer or cell lineage.

In naïve CD4^+^ T cells, features dysregulated during the first wave (∼age 40) overlapped substantially with Cluster 1 and were significantly enriched in cell cycle-related pathways (Fig. 3E, 3F and S7F-S7G). Conversely, molecular shifts during the second wave (∼age 70 year) largely coincided with Cluster 3, which exhibited a global upregulation trend and was significantly enriched in pathways associated with T cell activation (Fig. 3E, 3F and S7H).

### Deciphering the age-associated regulatory remodeling of transcribed CREs

Cis-regulatory elements (CREs) can be transcribed into stable or unstable RNAs; specifically, capped transcripts generated bidirectionally from active enhancers are termed enhancer RNAs (eRNAs)^33,34^. These transcribed CREs have recently emerged as robust proxies for enhancer activity, providing higher functional relevance than elements defined solely by histone marks or chromatin accessibility^10,34,35^. By integrating open chromatin regions from the CIMA scATAC-seq cohort with TSS signals derived from 5′ scRNA-seq within the same cell type, we classified ATAC peaks based on their transcriptional status and genomic proximity to promoters (Fig. 4A). Across the three most abundant cell types, the vast majority of proximal ATAC (pATAC) peaks were actively transcribed (Fig. 4B). The enrichment of H3K4me3, a canonical promoter hallmark^36^, at these transcribed proximal ATAC peaks in naïve CD4^+^ T cells underscores their identity as active transcription start sites (Fig. 4C, left). In contrast, only 21.6% (13,562/62,904) of distal ATAC (dATAC) peaks in naïve CD4+ T cells exhibited detectable TSS signals, compared to 71.1% (19,052/26,792) of proximal peaks (Fig. 4B). This trend, consistently observed in CD14^+^ monocytes and CD56^dim^ NK cells, indicates that distal elements generally possess lower transcriptional output than their proximal counterparts. Notably, transcribed distal ATAC peaks in naïve CD4^+^ T cells displayed higher densities of H3K27ac and H3K4me1 compared to their non-transcribed counterparts (Fig. 4C, right), suggesting that coupling RNA transcription with chromatin accessibility more effectively captures functional enhancer regions.

**Figure 4.**
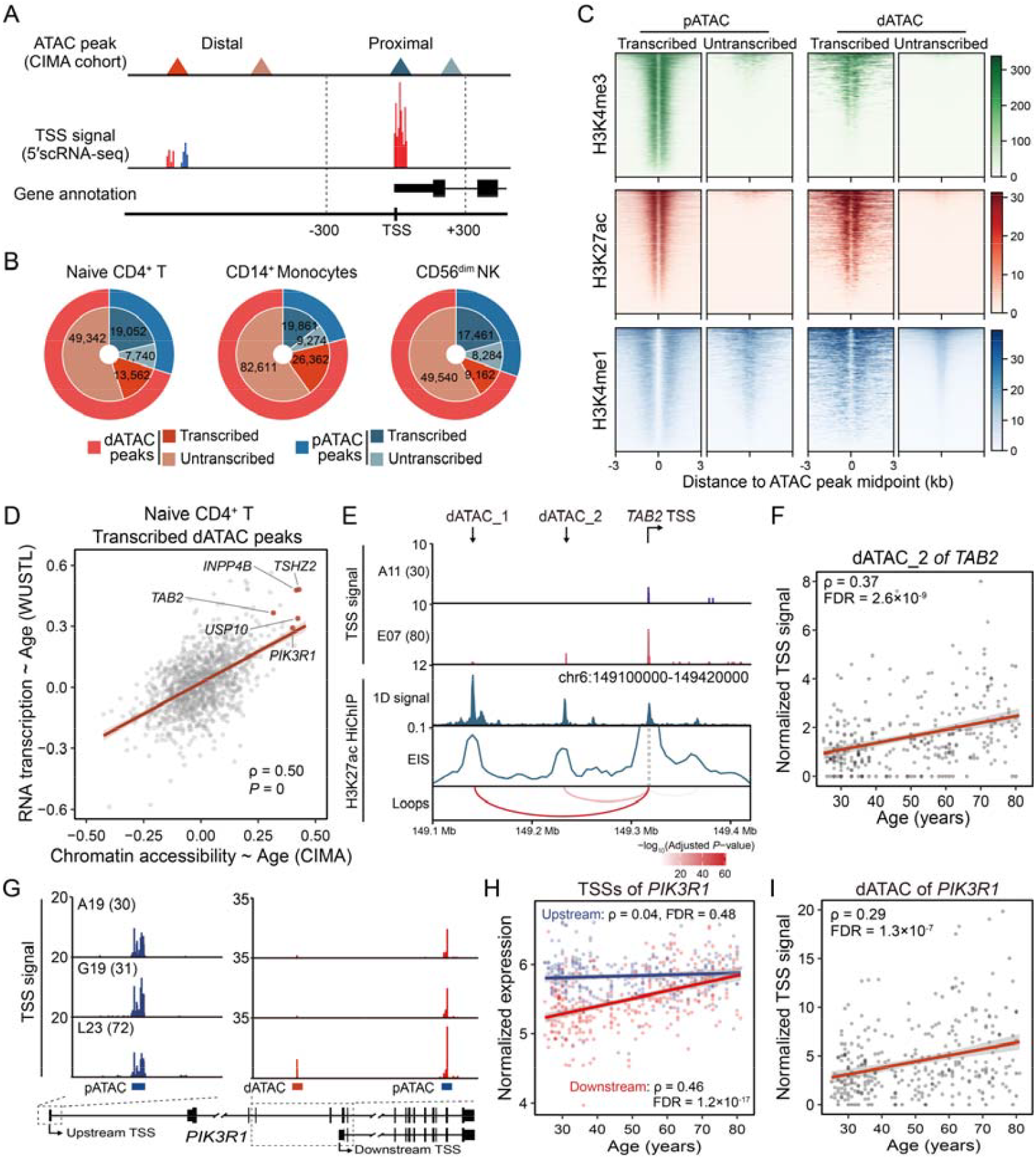
Dynamics of transcribed CREs during aging of CD4+ Naïve T cells. (A) Schematic illustrating the categorization of ATAC-seq peaks from the CIMA cohort based on TSS signal (5′ scRNA-seq) and genomic proximity to promoters. Peaks within ±300 bp of a transcript’s 5′ end were defined as proximal (pATAC), where those outside this window were considered distal (dATAC). (B) Pie chart showing the proportion of transcribed versus untranscribed ATAC peaks across three representative cell types in the CIMA cohort. (C) Heatmaps depicting normalized signal intensities for H3K4me3, H3K27ac and H3K4me1 in CD4+ T cells (ENCODE datasets) ±3kb around the midpoints of ATAC peaks from each category. (D) Scatter plot showing the Spearman correlation between chronological age and either chromatin accessibility (CIMA cohort, x-axis) or RNA expression (WUSTL cohort, y-axis) for transcribed dATAC peaks in CD4+ naïve T cells. (E) Integrative tracks illustrate normalized TSS density profiles, HiChIP-derived 1D H3K27ac enrichment and promoter-centric interaction profiles at the *TAB2* locus. Significant loop interactions are visualized and colored by their adjusted *P* values. (F) Spearman correlation between chronological age and RNA transcription at the peak dATAC_2 of *TAB2* in CD4+ naïve T cells. (G) Normalized TSS density profiles showing age-related shifts in RNA expression for three distinct ATAC-seq peaks within the *PIK3R1* locus in CD4+ naïve T cells. Transcript annotations are shown below. (H) Spearman correlation between chronological age and the expression levels of two distinct FE of *PIK3R1*. (I) Spearman correlation between chronological age and RNA transcription of the dATAC peak within *PIK3R1* in CD4+ naïve T cells.

Building upon our identification of transcribed CREs, we extended our analysis to evaluate how aging reshapes the regulatory coordination between RNA transcription and chromatin accessibility. At a global level, we identified a significant positive correlation between aging-related dynamics in chromatin accessibility and RNA transcription across all transcribed pATAC peaks, particularly in naïve CD4+ T cells (Spearman’s ρ = 0.2, *P* = 5.5×10^-87^, Fig. S7A). A representative instance is the pATAC peak at the *SOX4* promoter, where a significant decline in accessibility during aging mirrored a corresponding reduction in its transcriptional output in naïve CD4+ T cells (Fig. S4B and S7B). For transcribed dATAC peaks in naïve CD4+T cells, we identified a robust positive correlation between aging-associated shifts in chromatin accessibility and transcriptional activity (Spearman’s ρ = 0.5, Fig. 4D). Specifically, a distal CREs located within the first intron of *TSHZ2* exhibited concomitant increases in accessibility and RNA transcription during aging of naïve CD4+ T cells (Fig. S7C and S7D), coinciding with a significant age-associated elevation in overall *TSHZ2* mRNA levels (Fig. S7E). Notably, *TSHZ2* overexpression has linked to impaired T cell proliferation and a sustained decline in cell count^37^.

To investigate the regulatory impact of these transcribed CREs, we integrated H3K27ac HiChIP data from primary naïve CD4+ T cells to map long-range chromatin interactions^38^. For example, we identified two distal transcribed ATAC peaks upstream of the *TAB2* gene that physically interact with the *TAB2* promoter (Fig. 4E). Although both dATAC peaks exhibited increased accessibility with age (Fig. S7F), only dATAC_2 displayed a corresponding elevation in transcriptional activity (Fig. 4F and S7G). As a scaffold protein and ubiquitin sensor within the TAK1-TAB complex, TAB2 is pivotal for activating MAPK and NF-κB signaling cascades that drive inflammatory responses^39,40^. Given the observed upregulation of *TAB2* in naïve CD4+ T cells from older individuals (Fig. S7H), our findings suggest that age-associated distal CRE-mediated transcription regulation of *TAB2* may contribute to immune dysregulation.

The *PIK3R1* locus encompasses two distinct TSSs separated by over 60 kilobase (kb) (Fig. 4G). The upstream TSS gives rise to the full-length isoform (encoding p85α), which stabilizes p110 catalytic subunits and regulates cell proliferation in response to receptor tyrosine kinase phosphorylation^41,42^. Interestingly, our analysis revealed a bifurcated regulatory response during aging: while the upstream TSS remained stable, the downstream TSS exhibited heightened chromatin accessibility and significantly upregulated expression in naïve CD4+ T cells from older individuals, a trend consistently observed across both WUSTL and SHPD cohorts (Fig. 4H and S7I). This downstream TSS encodes a truncated isoform that lacks the regulatory SH3 and BH domains but retains the iSH2 domain required for p110 binding (Fig. S7F). Notably, a dATAC peak situated 5 kb upstream of this secondary TSS showed synchronized gains in both accessibility and transcriptional output with age (Fig. 4I and S7J). The strong expression correlation between this dATAC peak and the downstream TSS (Spearman’s ρ = 0.33, *P* = 1.6×10^-9^), standing in stark contrast to its negligible correlation with the upstream TSS (Spearman’s ρ = 0.14, *P* = 0.013), indicates that the age-associated dynamics of *PIK3R1* are driven by a specific enhancer-promoter unit rather than a global, locus-wide alteration.

### Age-associated reactivation of endogenous retrovirus reshapes T cell transcriptome

Transposable elements (TEs) constitute approximately half of the human genome and exert a profound influence on host gene-regulatory networks, emerging as central instructors of immune cell functions^12,43^. However, a comprehensive high-resolution analysis of aging-related TE expression dynamics within the immune system remains elusive. Here, we leveraged robust profiling of TSS signals to characterize the dynamics of TE expression across diverse immune cell populations during aging. Our analysis focused on retrotransposons (LINE, LTR, and SINE) and DNA transposons, owing to their significant regulatory potential^44^. By quantifying TSS-associated reads across subclasses, we identified millions of reads originating from TEs (Fig. S8A), providing sufficient resolution to interrogate aging-associated transcriptional shifts. Notably, we observed a striking positive linear correlation between LTR element expression and chronological age across diverse cell types in both cohorts (Fig. 5A). This age-related transcriptional reactivation was particularly pronounced in naïve CD4+ T and γδ T cells (Fig. 5B, S8B-C). Detailed examination of specific LTR families revealed that multiple endogenous retrovirus (ERVs), notably LTR32 and MLT2B3, exhibited elevated expression in naïve CD4+ T cells from older individuals (Fig. 5C). By comparing RNA expression of ERV copies between the older (65-81 years) and young (25-35 years) groups, we identified 29 copies that were significantly upregulated with age in naïve CD4+ T cells (Fig. 5D). Consistently, these transcriptionally active ERV copies displayed increased chromatin accessibility in naïve CD4+ T cells from older individuals (Fig. 5E), suggesting an epigenetic basis for their age-associated reactivation.

**Figure 5.**
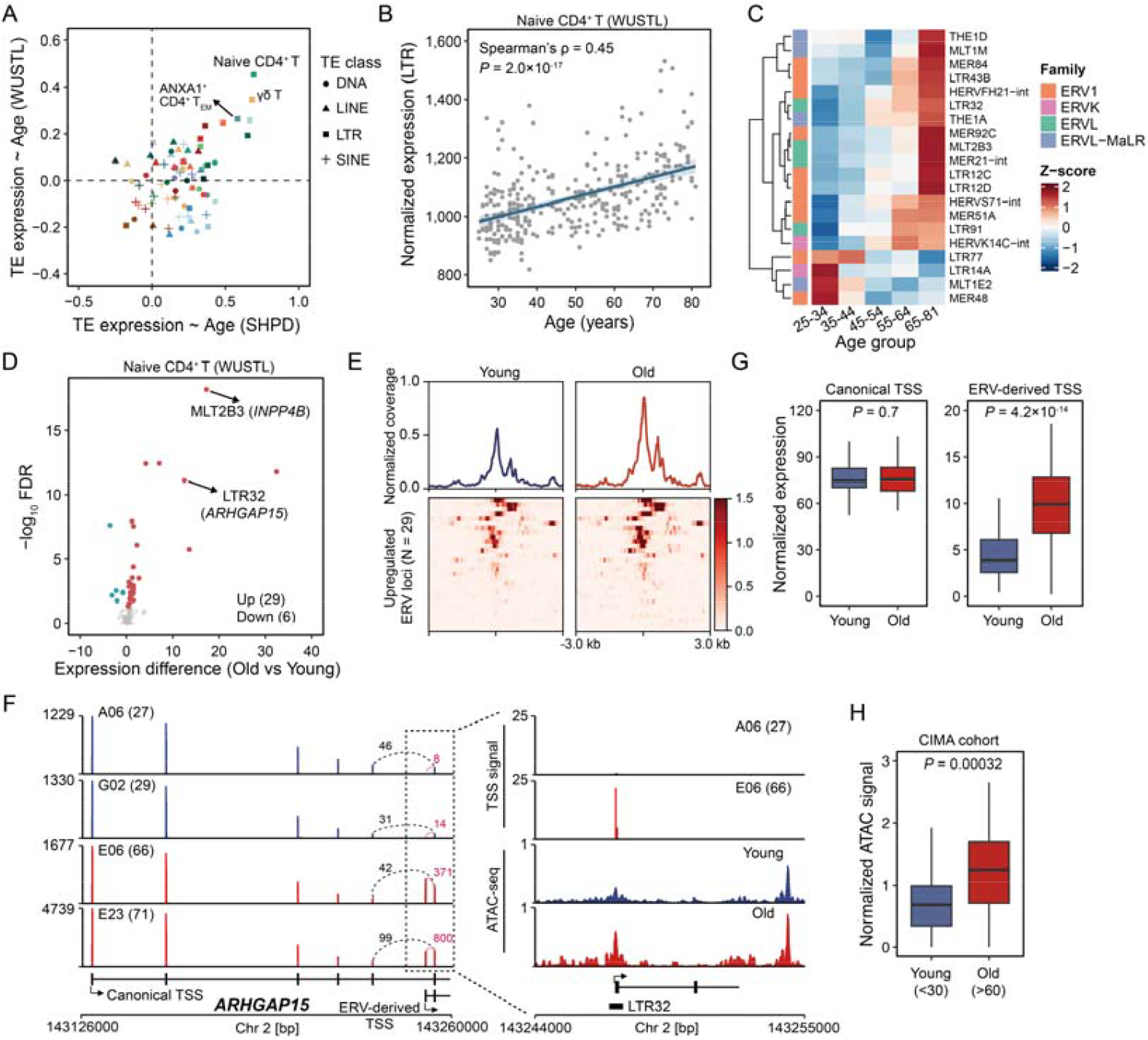
Age-related reactivation of ERVs in T cells. (A) Spearman correlation of total expression level from distinct classes of transposable elements with chronological age across cell types. Correlations for the SHPD cohort are shown on the x-axis and for the WUSTL cohort on the y-axis, with each point representing a TE class within a specific cell type. (B) Increased transcriptional activity of LTR elements with aging in CD4+ naïve T cells. Spearman correlation coefficient and *P* value are shown. (C) Age-related expression shifts of different LTR subfamilies in CD4+ naïve T cells. Five age groups of donors from the WUSTL cohort were shown. (D) Differential expression of individual LTR copies in CD4+ naïve T cells between old (65-81 years) and young (25-34 years) donors form the WUSTL cohort. Each point represents one LTR copy. (E) Chromatin accessibility profiles of isolated bulk CD4+ naïve T cells from young and old individuals, centered on LTR copies that show increased expression with age. (F) Left: Pseudobulk scRNA-seq tracks of CD4+ naïve T cells illustrating elevated expression of a short *ARHGAP15* isoform originating from an ERV-derived promoter in old individuals. Right: Normalized TSS and ATAC signals supporting the activation of ERV-derived short isoform. (G) Comparison of normalized expression levels of the canonical long isoform (left) and the ERV-derived short isoform (right) of *ARHGAP15* in CD4+ naïve T cells from young and old individuals in the WUSTL cohort. (H) Increased chromatin accessibility at the promoter of the short *ARHGAP15* isoform in CD4+ naïve T cells from old individuals in the CIMA cohort.

Intriguingly, among the top up-regulated ERV loci in naïve CD4+ T cells, we identified two acting as alternative promoters that drive the expression of chimeric transcripts (Fig. 5D). Specifically, an intronic MLT2B3 element mediated the transcription of an *INPP4B* short isoform (Fig. S8D-E), while an LTR32 element, situated with the fifth intron of *ARHGAP15*, initiated a truncated transcript (Fig. 5F). Analysis of TSS signals and splice junction reads confirmed that this LTR32-driven internal initiation yields a truncated *ARHGAP15* mRNA, whereas the canonical full-length transcripts remained non-differentially expressed between age groups in both the WUSTL and SHPD cohort (Fig. 5G and S8F). Consistent with these transcriptional shifts, ATAC-seq data revealed significantly increased chromatin accessibility at this ERV locus in older individuals (Fig. 5H and S8G). ARHGAP15 is known to promote the hydrolysis and inactivation of the Rac1 GTPase^45^, a pivotal signaling hub that couples T cell receptor (TCR) activation to the induction of *IL2* and *IL2RA*, thereby driving T cell expansion^46^. Reanalysis of a published genome-scale T cell CRISPR screening dataset^47^ further identified *ARHGAP15* as a top hit, whose overexpression reduces IL-2 cytokine production in CD4+ T cells (Fig. S8H). Notably, protein architecture analysis confirms that the LTR32-initiated isoforms lack the N-terminal PH domain while preserving the intact catalytic RhoGAP domain (Fig. S8I). The functional impact of these truncated transcripts on naïve CD4+ T cells warrant further experimental validation. In summary, our findings demonstrate that epigenetic reactivation of specific ERVs serves as a molecular feature of human T cell aging.

### Age-related increase in EDA expression contributes to the functional decline of naïve CD4+ T cell responses in older individuals

Among the genes exhibiting age-related expression dynamics in naïve CD4+ T cells, *EDA* stood out with one of the strongest correlations with age across both the WUSTL and SHPD cohort (Fig. 2D, 6A, 6B and S9A). This age-associated linear increase in *EDA* expression was consistently validated in two additional independent cohorts: the Immunology of Aging and CIMA cohorts (Fig. 6C and S9B). Furthermore, bulk ATAC-seq profiling of isolated naïve CD4+ T cells revealed significantly enhanced chromatin accessibility at the *EDA* promoter region in older individuals (Fig. 6A and S9C). This trend was recapitulated in the scATAC-seq data from the CIMA cohort (Fig. 6D), indicating the age-associated epigenetic activation of the *EDA* locus in naïve CD4+ T cells.

**Figure 6.**
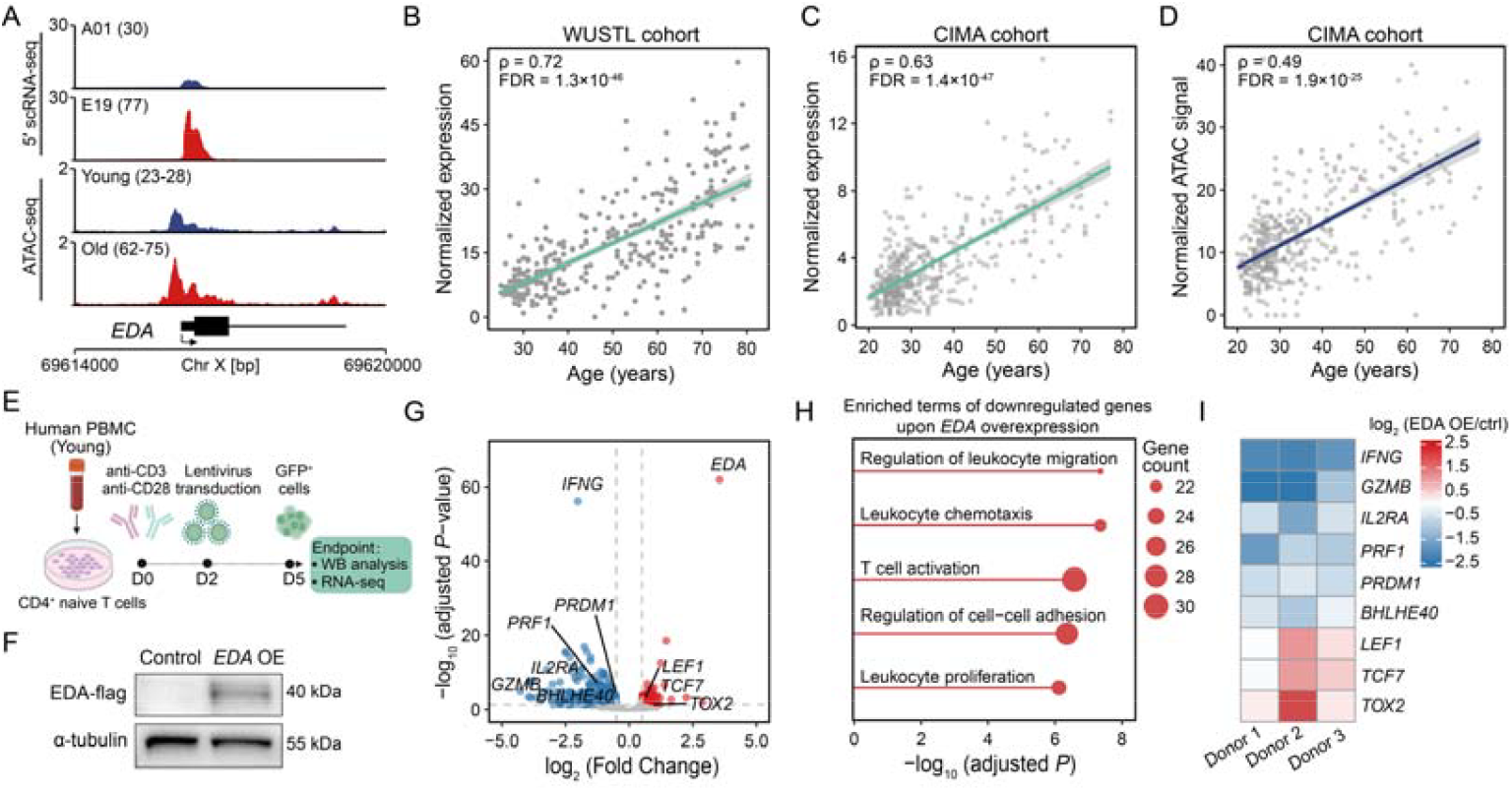
EDA upregulation contributes to functional decline of CD4^+^ naïve T cell. (A) Normalized 5′ scRNA-seq (top) and ATAC-seq (bottom) tracks showing elevated *EDA* expression and increased chromatin accessibility at its promoter region in CD4+ naïve T cell from older compared to younger individuals. (B, C) Spearman correlation between chronological age and *EDA* expression in CD4+ naïve T cells from the WUSTL (B) and CIMA (C) cohorts. (D) Age-associated increase in chromatin accessibility at the *EDA* promoter in CD4+ naïve T cells from the CIMA cohort. (E) Schematic for in vitro experiment of *EDA* overexpression in CD4+ naïve T cells. The experiments were performed three times independently using PBMCs from different donors (aged 20-25 years). (F) Western blot confirming the successful overexpression of *EDA* protein in primary CD4+ naïve T cells isolated from a healthy young donor. (G) Volcano plot displaying differentially expressed genes following *EDA* overexpression in primary CD4+ naïve T cells (n = 3 biological replicates from independent donors). Red and blue dots represent significantly up- and down-regulation genes, respectively (|log_2_ fold change| > 0.5, adjusted *P* < 0.05). (H) Gene Ontology enrichment analysis of genes significantly downregulated upon *EDA* overexpression. (I) Heatmap illustrating the differential expression of representative genes upon *EDA* overexpression.

To investigate the functional consequences of EDA overexpression, we isolated primary naïve CD4+ T cells from three healthy young individuals and performed lentiviral transduction for control or EDA overexpression after in vitro stimulation. Successfully transduced GFP+ T cells were sorted for bulk RNA-seq to characterize the transcriptional dynamics (Fig. 6E). Increased EDA protein expression after transduction was confirmed by Western blotting (Fig. 6F). We observed widespread gene expression changes upon *EDA* overexpression, identifying 364 significantly downregulated and 102 significantly upregulated genes (Fig. 6G). Notably, genes downregulated upon EDA overexpression were significantly enriched in pathways involved in T cell activation and effector function, such as “T cell activation”, “Leukocyte chemotaxis”, and “leukocyte proliferation” (Fig. 6H). In line with these findings, effector genes (e.g., *IFNG, GZMB, IL2RA, PRF1*, and *PRDM1*) displayed maximal reduction upon *EDA* overexpression (Fig. 6I), whereas genes characteristic of low activation or exhausted states (e.g., *LEF1, TCF7*, and *TOX2*) were most highly expressed in *EDA* overexpressed group (Fig. 6I). Together, these data suggest that increased *EDA* expression contributes to impaired T cell responses, potentially accounting for age-related immune deficiency.

## Discussion

In this study, we employed 5′ scRNA-seq to precisely resolve TSSs and splice junctions across peripheral immune cell populations at single-cell resolution. Moving beyond conventional gene-level expression analysis, we provide the first comprehensive demonstration of the widespread dynamics of alternative TSS usage and splice site selection during immune aging. Leveraging the unique advantages of 5′ scRNA-seq for TSS profiling, we further quantified the transcriptional activity of distal cis-regulatory elements and transposable elements, uncovering a previously unappreciated layer of age-related transcriptomic remodeling.

Recent multi-omics profiling has uncovered non-linear trajectories during human aging, identifying pivotal transition points throughout the lifespan^9,30-32^. For instance, longitudinal analysis of a deep multi-omics cohort (108 individuals aged 25-75□years) revealed two waves of molecular dysregulation at approximately 44 and 60□years of age^30^. A large-scale scRNA-seq study (230 donors aged 20 to 84 years) identified mid-life (∼40 years) as a critical inflection point for immune remodeling^9^. Utilizing multi-layer transcriptomic features of peripheral immune cells from the WUSTL cohort (n = 317, aged 25 to 85 years), we identified two prominent, global waves of transcriptomic remodeling centered at approximately ages 40 and 70 (Fig. 3D). The emergence of the first wave (∼40 years) aligns with recent reports^9,32^, underscoring the robustness of this mid-life transition. To further validate these nonlinear dynamics, we examined gene-level expression shifts in two additional cohorts. The Immunology of Aging cohort (234 donors aged 40 to >90 years) exhibited a similar bimodal pattern, with waves peaking at approximately ages 53 and 85 (Fig. S10A, B). In contrast, the CIMA cohort, which primarily comprises younger individuals (72% under 40 years), displayed a single prominent wave centered around age 50 (Fig. S10C, D). These variations in transcriptomic trajectories likely stem from a combination of biological and technical factors, including diverse geographic and genetic backgrounds^26,27^, differing age distributions across cohorts, and the use of distinct scRNA-seq platforms.

Consistent with prior reports highlighting the susceptibility of T cell subsets to aging^6,9^, we observed that these populations exhibit the most extensive transcriptomic remodeling during aging (Fig. 2A and S6A). Notably, we observed a significant age-associated contraction of the naïve CD8+ T cell compartment, whereas the naïve CD4+ T cell population remained relatively stable across the lifespan (Fig. S10E). This stability, coupled with the profound transcriptomic shifts detected, identified naïve CD4+ T cells as an appropriate population for characterizing age-related features. In contrast, the diminished cell counts of naïve CD8+ T cells in older individuals led to a reduction of total sequenced reads available for pseudobulk analysis (Fig. S10F), potentially limiting the sensitivity of TSS quantification and splice junction detection in this subset. Further studies employing increased cell number per donor (particularly for older individuals) or greater sequencing depth will be essential to further refine isoform-level transcriptomic landscapes in these contracting cell populations.

Vaccine efficacy against many pathogens in older adults is attenuated due to the aging of naïve CD4+ T cells, the central orchestrator of vaccine responses^48^. Age-associated impairments in TCR signaling of naïve CD4+ T cells have been reported, in part driven by reduced expression of miR-181a and the subsequent accumulation of several phosphatases, including dual-specific phosphatase 6 (DUSP6)^49^. Here, we demonstrate that *EDA* expression increased progressively with age, and its over induction in in vitro-stimulated in naïve CD4^+^ T cells after activation markedly suppresses gene networks essential for T cell activation and effector function. These findings establish *EDA* as a potentially complementary axis contributing to age-associated T cell dysfunction. While we observed increased chromatin accessibility at the *EDA* promoter with age, the upstream regulators driving its upregulation, as well as the downstream biochemical cascades through which *EDA* regulates T cell activation, remain to be fully elucidated.

In summary, our findings reveal that immune aging is characterized by multi-layered transcriptomic shifts that extend far beyond simple changes in gene-level expression abundance. These high-resolution insights provide a valuable resource for understanding the regulatory programs that govern the functional decline of the human immune system over time.

## Methods

### Data collection

We assembled 5′ scRNA-seq data from two independent cohort. The first dataset (WUSTL cohort) comprises 317 samples from 166 healthy individuals (aged 25-81 years), as established by the Washington University School of Medicine, Saint Louis (available via Synapse: syn49637038)^7^. The second dataset (SHPD cohort) consists of 61 healthy individuals from the Shanghai Pudong Cohort (0 to >90 years, available at the Genome Sequence Archive for Human under accession number HRA009014)^8^. To complement these, processed scRNA-seq and scATAC-seq data from the CIMA cohort were obtained from (https://db.cngb.org/trueblood/cima/)^3^. Furthermore, we incorporated bulk ATAC-seq (GEO: GSE179593)^50^, bulk RNA-seq (SRA: PRJNA757466)^51^ and bulk H3K27ac HiChIP data (GEO: GSE101498)^38^ of primary naive CD4+ T cells isolated from different donors. We retrieved ChIP-seq datasets for key histone modifications in CD4+ T cells from the ENCODE database (https://encodeproject.org), including H3K27ac (ENCFF018YIU), H3K4me1 (ENCFF860QIV), and H3K4me3 (ENCFF610SPR).

### Isolation and culture of human primary CD4+ naïve T cells

Human PBMCs were obtained healthy blood donors recruited at Huadong Hospital (Shanghai, China). The study was conducted in accordance with the principles of the Declaration of Helsinki and approved by the Huadong Hospital Institutional Review Board (protocol #2023K189). Informed written consent was obtained from all participants prior to their inclusion in the study. Primary CD4+ naïve T cells were isolated using the EasySep™ Human Naive CD4+ T Cell Isolation Kit (STEMCELL Technologies) following the manufacturer’s protocol. The purity of the isolated populations was verified by flow cytometry. Purified cells were cultured in ImmunoCult™-XF T Cell Expansion Medium supplemented with 100 IU/ml recombinant human IL-2 (T&L Biotechnology). For T cell activation, cells were stimulated with ImmunoCult™ Human CD3/CD28 T Cell Activator (STEMCELL Technologies).

### Lentivirus production and transduction

The human *EDA* coding sequence was cloned into HBLV-3Flag-ZSGreen-PURO lentiviral vector, with the HBTLV-ZsGreen-PURO empty vector serving as a negative control (Hanbio Biotechnology). To produce lentiviral particles, HEK293T cells were co-transfected with the respective lentiviral vectors and the packaging plasmids psPAX2 (Addgene #12260) and pMD2.G (Addgene #12259) using Lipofiter™ (Hanbio Biotechnology). Viral supernatants were harvested at 48 and 72 h post-transfection, pooled, clarified by centrifugation (2,000 × g, 10 min, 4°C), and further concentrated via ultracentrifugation (82,700 × g, 120 min, 4°C). The resulting viral pellets were resuspended in chilled medium and stored at −80°C. Primary CD4^+^ naïve T cells were pre-activated for 48 h prior to transduction. Cells were then infected at a MOI of 150 in the presence of 6.5 μg/ml polybrene and 100 U/ml IL-2. Two days post-transduction, ZsGreen-positive cells were isolated via fluorescence-activated cell sorting and subsequently subjected to RNA-seq analysis.

### Western blotting

Cells were lysed in RIPA buffer (Thermo Fisher Scientific) supplemented with protease and phosphatase inhibitors (Beyotime) for 20 minutes on ice. Equal amounts of proteins were resolved on 8-16% SDS-PAGE gels (GenScript) before being transferred onto PVDF membranes (Vazyme). After blocking with 5% non-fat milk in TBST for 1 h at room temperature, the membranes were incubated overnight at 4°C with the following antibodies: anti-FLAG (F1804, Sigma-Aldrich) and HRP-conjugated anti-α-tubulin antibodies (66031, Proteintech) as a loading control. After washing, the membranes were incubated with HRP-conjugated secondary antibodies (Abmart) for 1 h. Protein bands were visualized using ECL substrate and captured with a chemiluminescence imaging system (Tanon).

