## Supplemental Figures for "Multi-layer transcriptomic characterization of age-related immune dynamics"

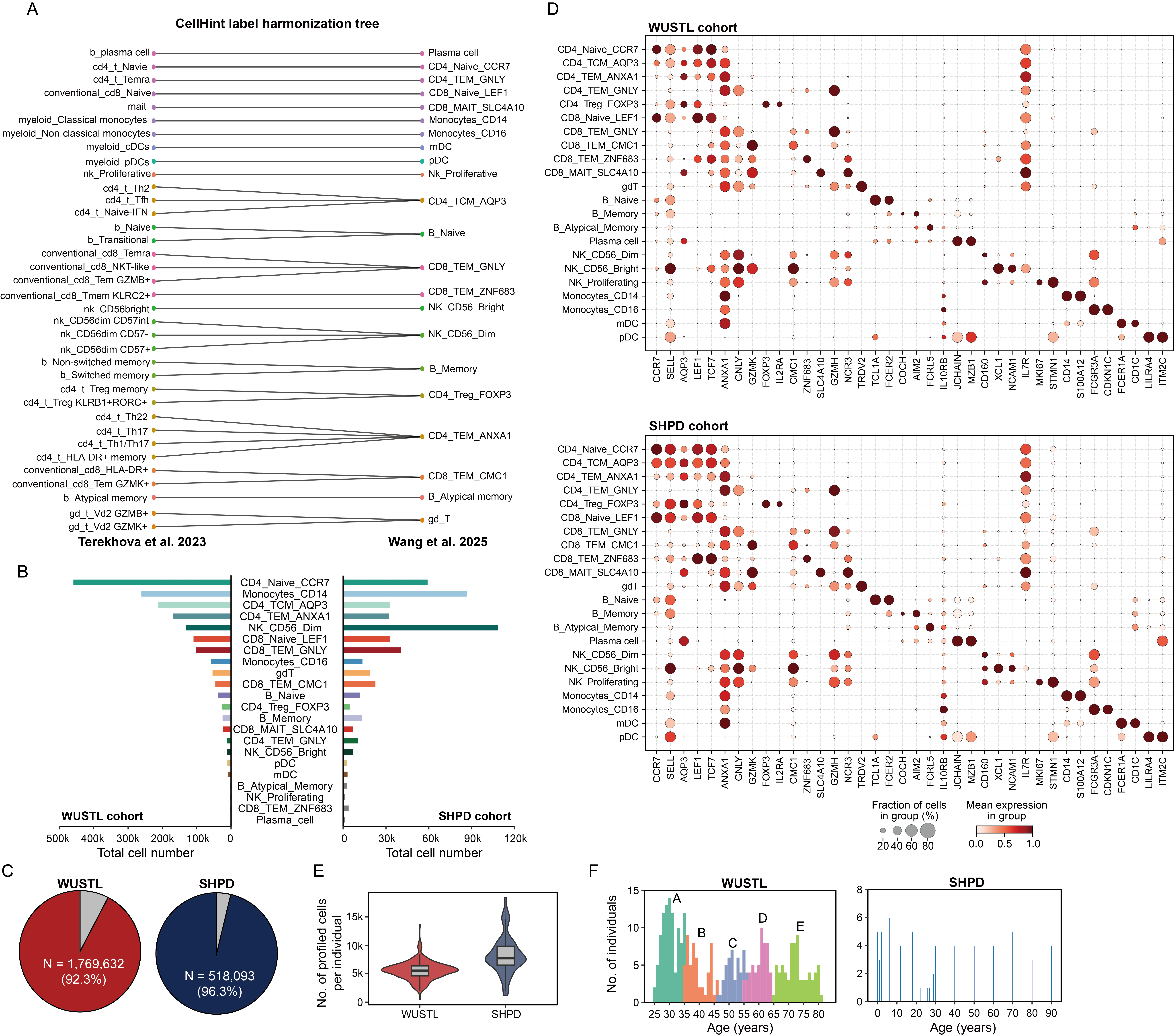

**Supplementary figure 1. Integration and overview of PBMC scRNA-seq data from two cohorts.**

(A) Harmonization graph illustrating cell-type relationships across two PBMC scRNA-seq datasets. Cell-type labels are colored according to their low-hierarchy alignments. The lines in the graph reflect the connections among transcriptionally similar cell types, independent of the original annotations. (B) Distribution of cell-type numbers across cohorts after harmonization. (C) Pie charts showing the number and proportion of all profiled cells retained after harmonization by CellHint. (D) Dot plots showing the expression levels of canonical marker genes across immune cell subtypes in the WUSTL (top) and SHPD (bottom) cohorts. Dot color represents the average scaled expression, and dot size indicates the fraction of cells expressing the gene within each subtype. (E) The number of retained cells per individual. (F) Age distribution across both cohorts.

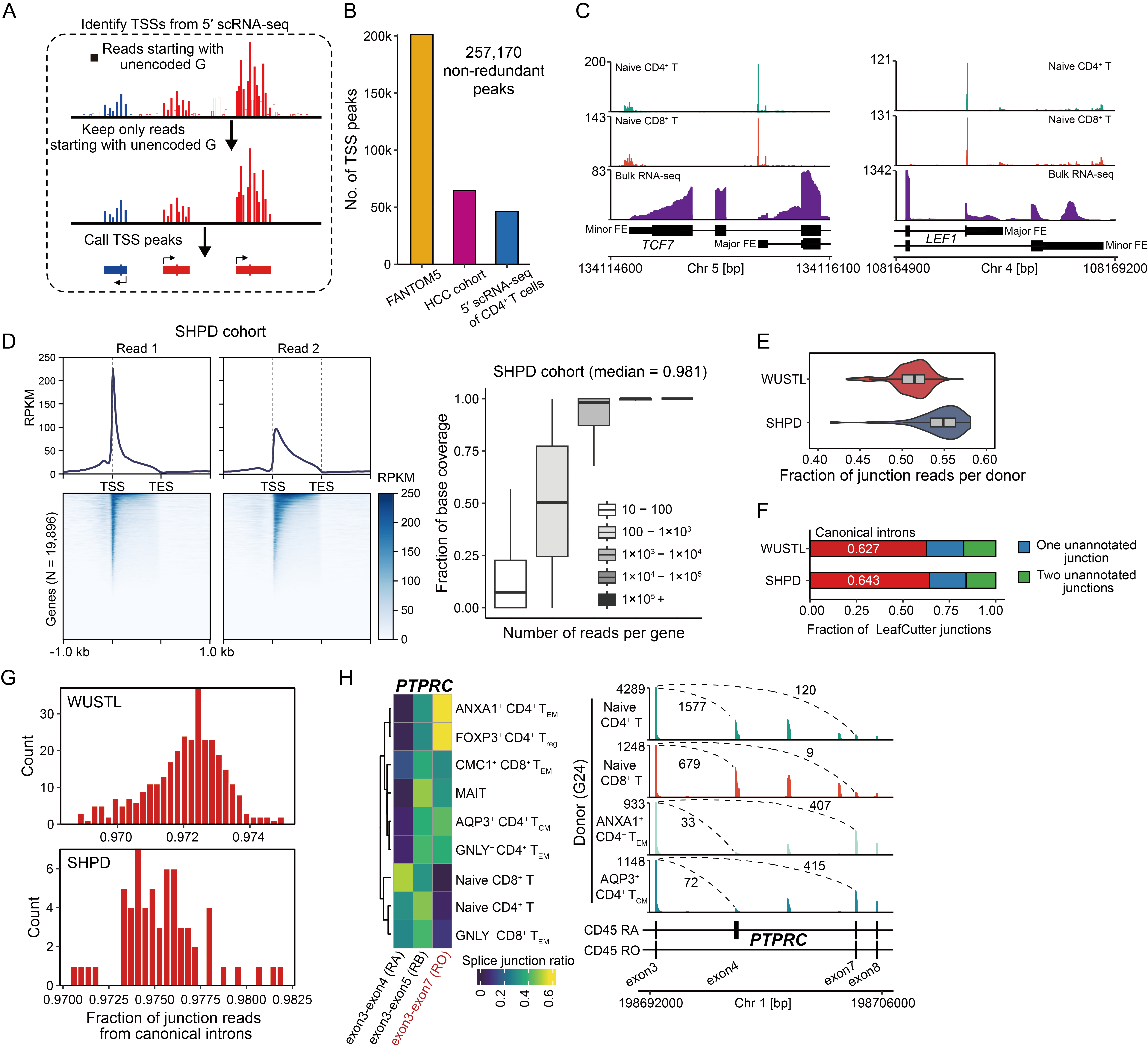

**Supplementary figure 2. Quality control of TSSs and splice junctions identified from 5' scRNA-seq.**

(A) Schematic overview of the ReapTEC approach. Only reads (read 1) starting with an unencoded “G” were retained for downstream analyses. This prefiltering step ensured high-confidence TSS peak calling and accurate TSS expression quantification. (B) The number of TSS peaks collected from previous studies. (C) TSS density profiles illustrating multiple detected TSS peaks in well-known genes with alternative promoters, including *TCF7* (left) and *LEF1* (right). (D) Left: Profile plot and heatmap showing the distribution of read 1 and read 2 across gene bodies in scRNA-seq data from the SHPD cohort. Right: Boxplot displaying the fraction of base coverage per gene across different read count bins. (E) Proportion of splice junction reads among all reads per individual in two cohorts. (F) Fraction of splice junctions identified by LeafCutter that are present in the GENCODE (v44) transcript annotation. (G) Fraction of splice junction reads that annotated by GENCODE (v44) transcript annotation per individual in two cohorts. (H) Left: Heatmap displaying the splicing junction ratios of *PTPRC* across T cell subsets, illustrating that mRNA expression of the CD45RO isoform (highlighted in red) was the lowest in naive T cells and increased in memory T cells. Right: Pseudo-bulk scRNA-seq tracks demonstrating alternative junction usage of *PTPRC* across T cell subtypes from donor A24 (tube id: G24).

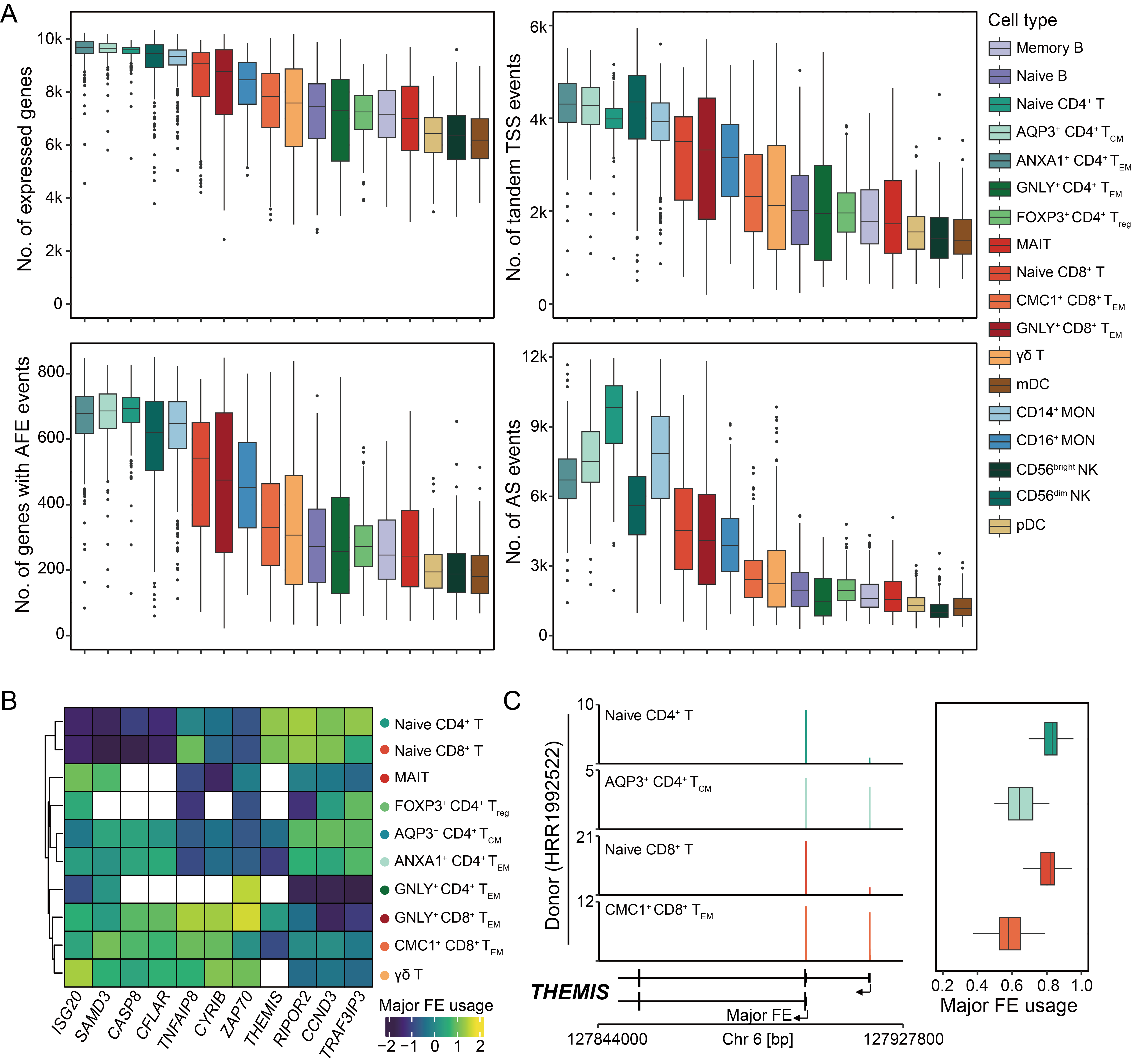

**Supplementary figure 3. Overview of detected transcriptomic features across cell types.**

(A) Number of transcriptomic features captured across four regulatory layers in pseudo-bulk samples across 18 cell types. (B) Heatmap of major FE usage for representative genes across T cell subsets. (C) Left: TSS read density profiles illustrating differential FE usage of *THEMIS* across pseudobulk T cell subsets from donor HRR1992522 (SHPD cohort). Right: Boxplot comparing the major FE usage of *THEMIS* across T cell subsets.

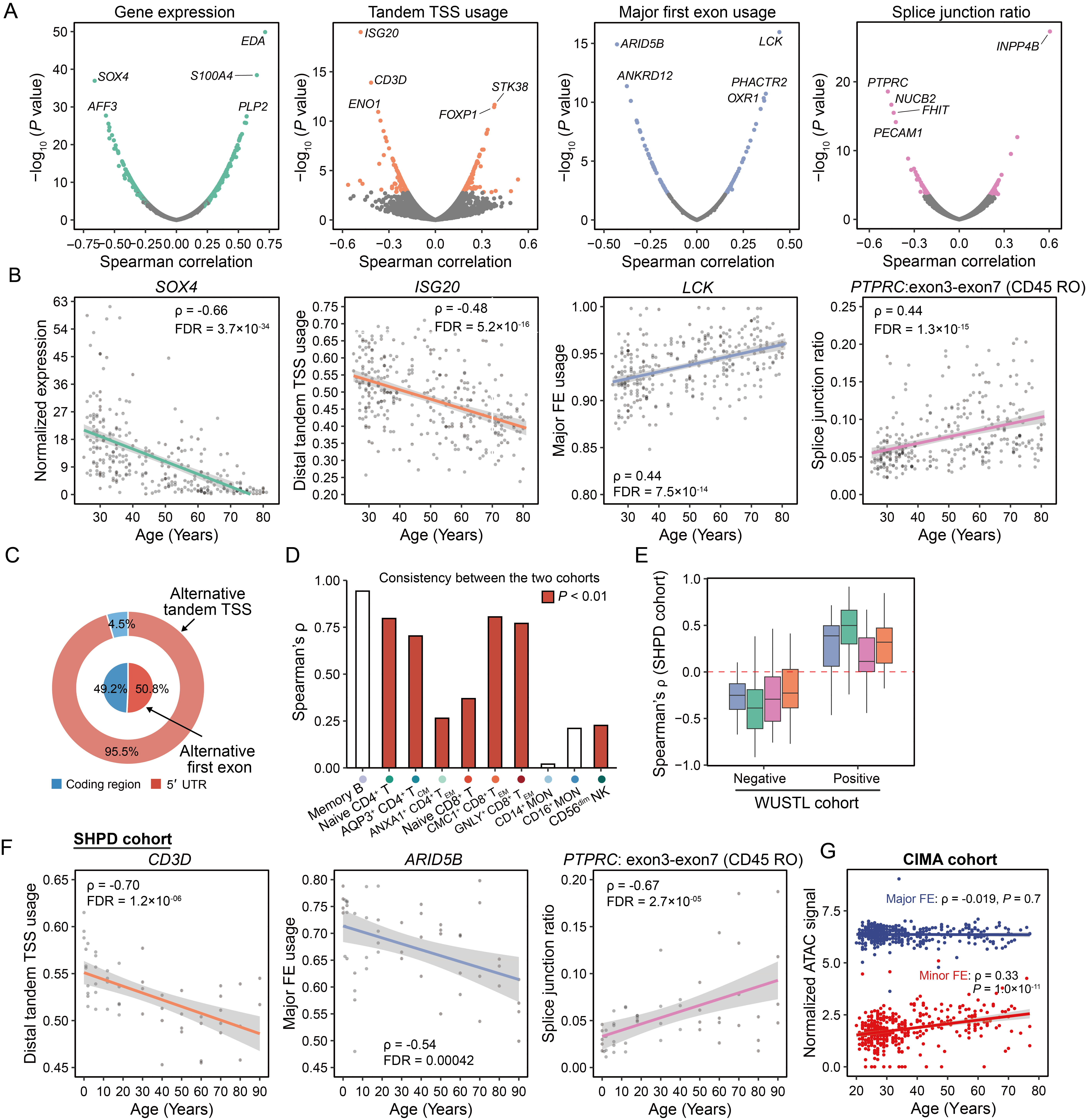

**Supplementary figure 4. Transcriptomic features showing linear changing with aging.**

(A) Scatter plot showing the Spearman correlation between different classes of transcriptomic features and chronological age detected in the CD4+ Naïve T cells. Significantly changed features were colored and the top five significantly correlated features were indicated. (B) Scatterplot displaying the correlation between age and four transcriptomic features in CD4+ naïve T cells (WUSTL cohort): overall *SOX4* expression, distal tandem TSS usage of *ISG20*, major FE usage of *LCK*, and the splice junction ratio (exon3-exon7, CD45RO) of *PTPRC*. (C) The effect of age-related dynamics in alternative TSS usage on gene product. (D) Correlation between the Spearman correlation coefficients calculated in the WUSTL cohort and SHPD cohort for all features exhibiting significant linear changes in the WUSTL cohort. (E) Distribution of Spearman correlation coefficients in the SHPD cohort for features that exhibited significant positive or negative linear changes in the WUSTL cohort. (F) Association between the distal tandem TSS usage of CD3D (left), the major FE usage of *ARID5B* (middle) and the splice junction ratio of *PTPRC* (right) with aging in CD4+ naïve T cells from the SHPD cohort. (G) Association between the chromatin accessibility of two FE region of *ARID5B* with aging in CD4+ naïve T cells from the CIMA cohort.

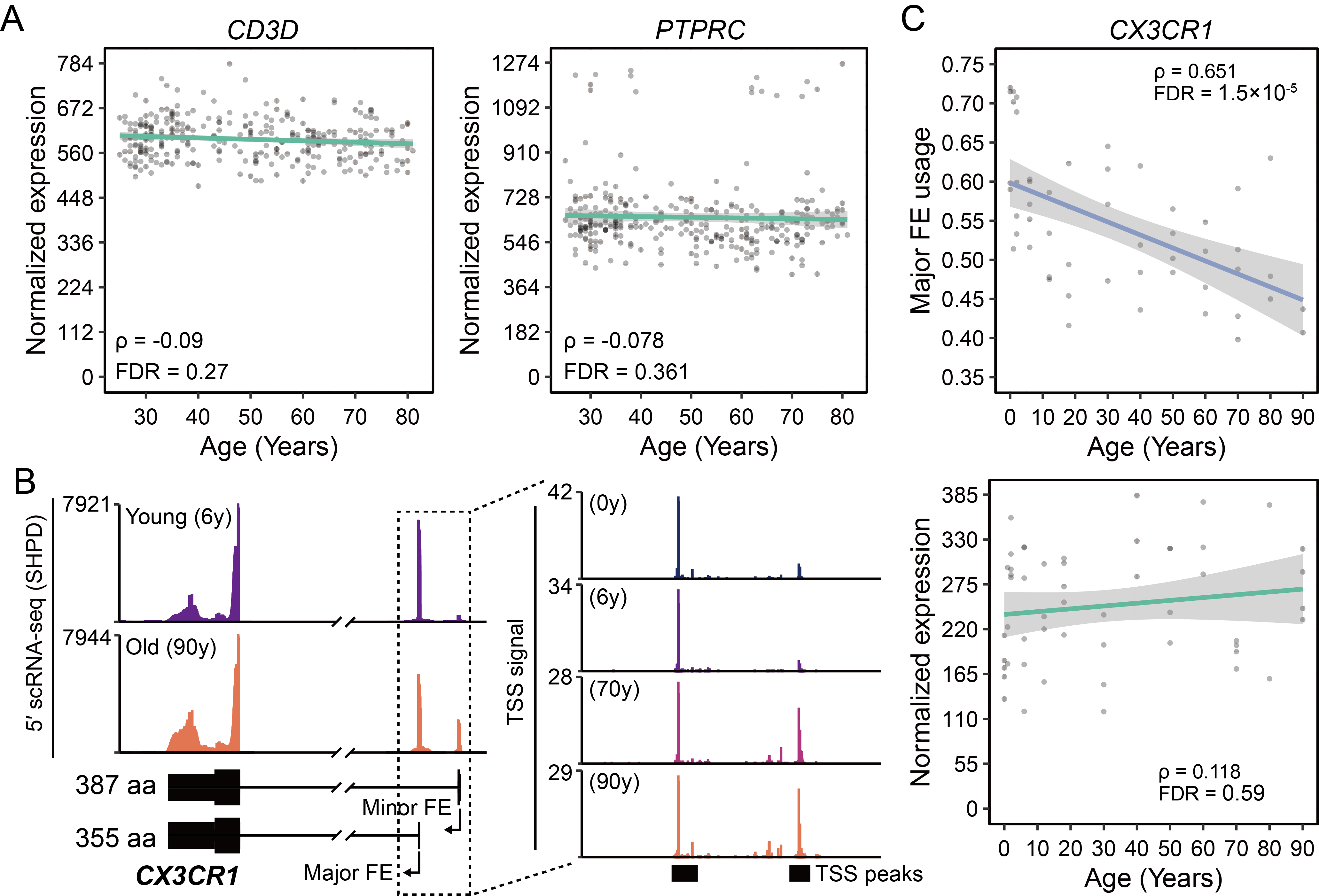

**Supplementary figure 5. Decoupling of age-related shifts in gene-level and isoform-level expression.**

(A) Association between the overall expression of *CD3D* (left) and *PTPRC* (right) with aging in CD4+ naïve T cells from the WUSTL cohort. (B) Pseudobulk scRNA-seq tracks illustrating the differential usage of *CX3CR1* alternative first exons in CD56^dim^ NK cells from donor with different age (SHPD cohort). The TSS signals originating from the two distinct first exons in four representative donors are highlighted. (C) Scatter plots showing the Spearman correlation between major first exon usage (top) and overall expression (bottom) of *CX3CR1* with chronological age in CD56^dim^ NK cells in the SHPD cohort.

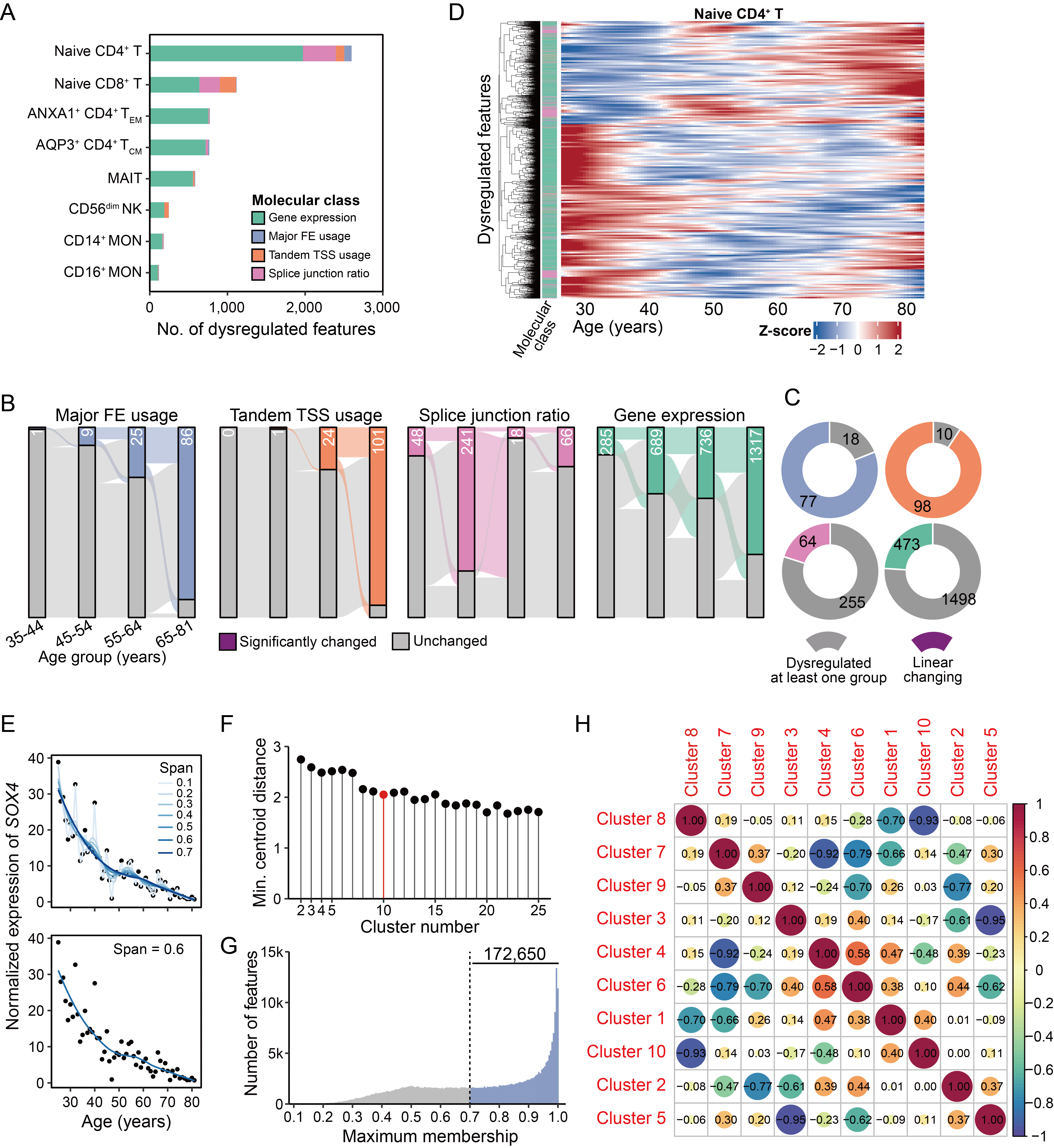

**Supplementary figure 6. Identification of nonlinearly changing transcriptomic features during aging using LOESS smoothing and fuzzy clustering.**

(A) Number of transcriptomic features significantly dysregulated in at least one age group among the top eight immune cell subtypes in the WUSTL cohort. (B) Sankey diagram illustrating the quantitative flow of significantly altered molecular features across distinct classes in CD4^+^ Naïve T cells, stratified by different age ranges compared to the baseline group (25-35 years old). (C) Donut charts depicting the proportion of linearly changing molecular features among those dysregulated in at least one age range in CD4^+^ Naïve T cells. (D) Heatmap depicting nonlinear age-associated changing of all dysregulated features in CD4+ Naïve T cells. (E) Top: LOESS smoothing curves of *SOX4* expression during aging using span parameters ranging from 0.1 to 0.7 (increment = 0.1). Bottom: LOESS smoothing of *SOX4* expression with the optimal span of 0.6. (F) Determination of the optimal number of clusters. (G) Distribution of maximum membership values across all molecular features; those with a maximum membership exceeding 0.7 were retained for downstream analysis. (H) Correlation matrix among the 10 identified clusters for all retained features.

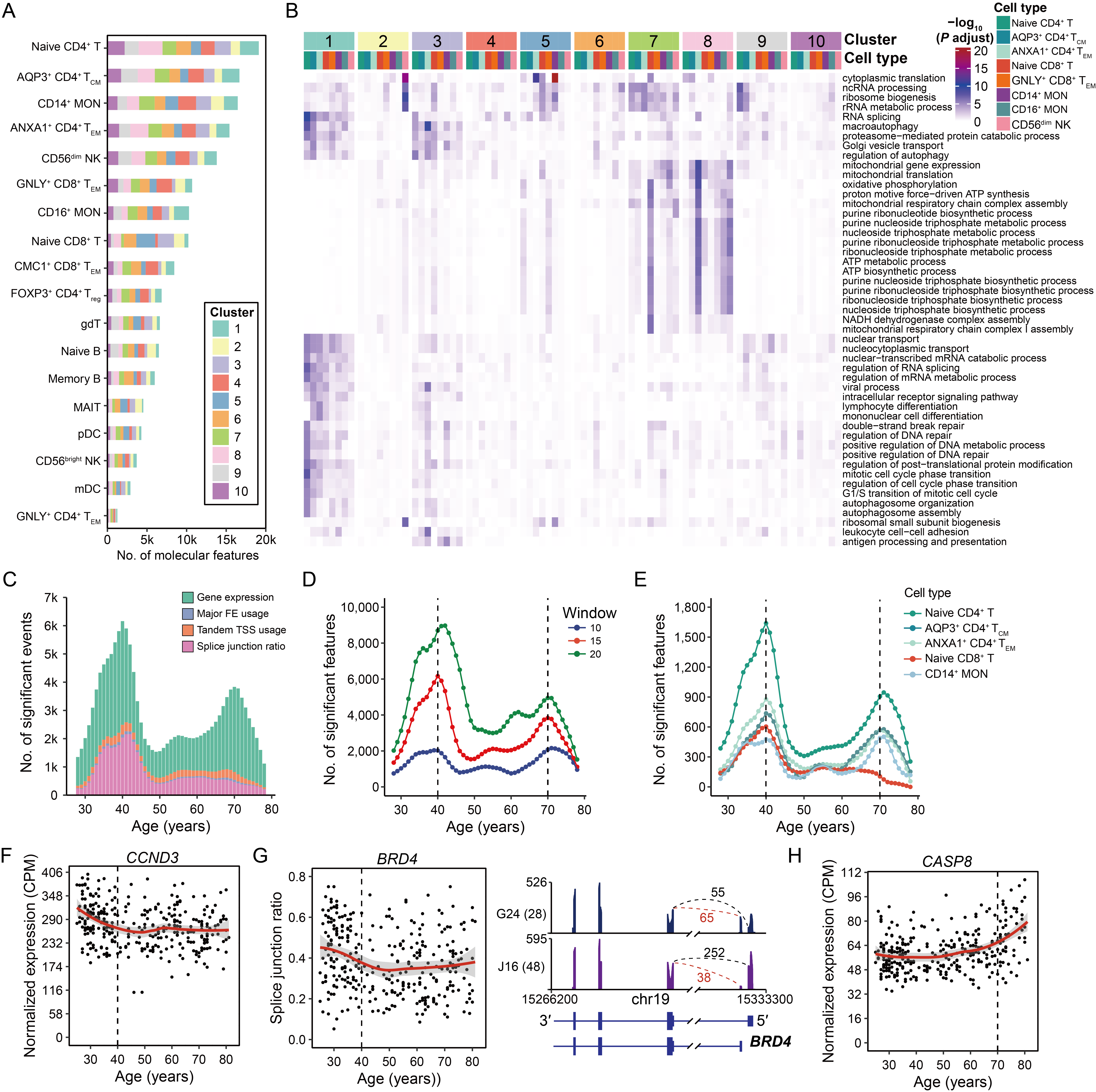

**Supplementary figure 7. Nonlinear dynamics of immune cells during aging in the WUSTL cohort.**

(A) Distribution of molecular features from each immune cell subtype across identified clusters. (B) Heatmap showing the GO enrichment for molecular features within each of the 10 clusters. (C) Number of significantly changed transcriptomic features, colored by distinct layers of gene regulation, during aging. (D) The same waves were detected using different window cutoffs. (E) Number of significantly changed transcriptomic features during aging across the five cell subtypes showing the most abundant changes. (F) Scatter plot showing the dynamics of *CCND3* expression in CD4+ naïve T cells during aging. (G) Left: Scatter plot showing the dynamics of *BRD4* alternative splicing in CD4+ naïve T cells during aging. Right: scRNA-seq tracks showing the alternative splicing of *BRD4* in donors with different ages. (H) Scatter plot showing the dynamics of *CASP8* expression in CD4+ naïve T cells during aging.

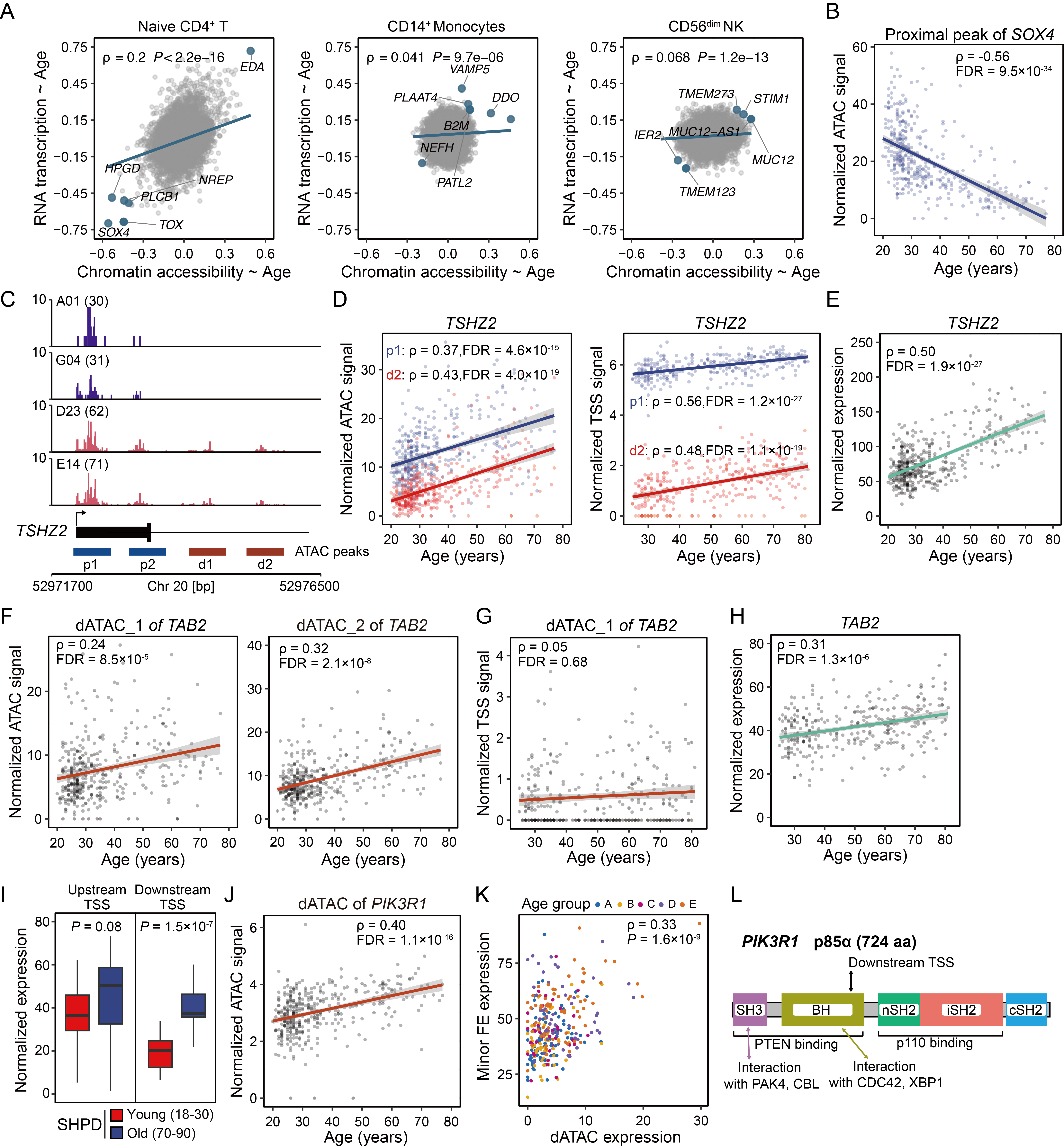

**Supplementary figure 8. Age-associated dynamics of transcribed cis-regulatory elements (CREs).** (A) Scatter plot showing the Spearman correlation with age for transcribed proximal ATAC peaks across CD4+ naïve T cells (left), CD14+ monocytes (middle), and CD56^dim^ NK cells (right). Data compare chromatin accessibility (CIMA cohort, x-axis) and RNA expression (WUSTL cohort, y-axis). (B) Age-dependent decrease in chromatin accessibility at the *SOX4* promoter region in CD4+ naïve T cells. (C) Normalized TSS signal illustrating the age-associated dynamics of transcribed CREs associated with *TSHZ2* in CD4+ naïve T cells. (D) Progressive increase in chromatin accessibility (left) and RNA expression (right) of the *TSHZ2* proximal peak (p1, red) and distal peak (d1, bule) during aging in CD4+ naïve T cells. (E) Upregulation of *TSHZ2* expression with aging in CD4+ naïve T cells. (F) Increased chromatin accessibility of the two distal peaks of *TAB2* during aging in CD4+ naïve T cells. (G) The correlation of RNA expression of the peak dATAC_1 of TAB2 with age in CD4+ naïve T cells. (H) Upregulation of *TAB2* expression with aging in CD4+ naïve T cells. (I) Boxplots comparing the expression of upstream TSS (left) and downstream TSS (right) of *PIK3R1* between young and old individuals in the SHPD cohort. (J) Increased chromatin accessibility of the *PIK3R1* distal ATAC peak during aging in CD4+ naïve T cells. (K) The correlation between RNA expression of distal ATAC peak and downstream FE in CD4+ naïve T cells. (L) Protein domain architecture of p85α encoded by *PIK3R1* full-length transcript. The downstream TSS generating N-terminally truncated transcripts was indicated.

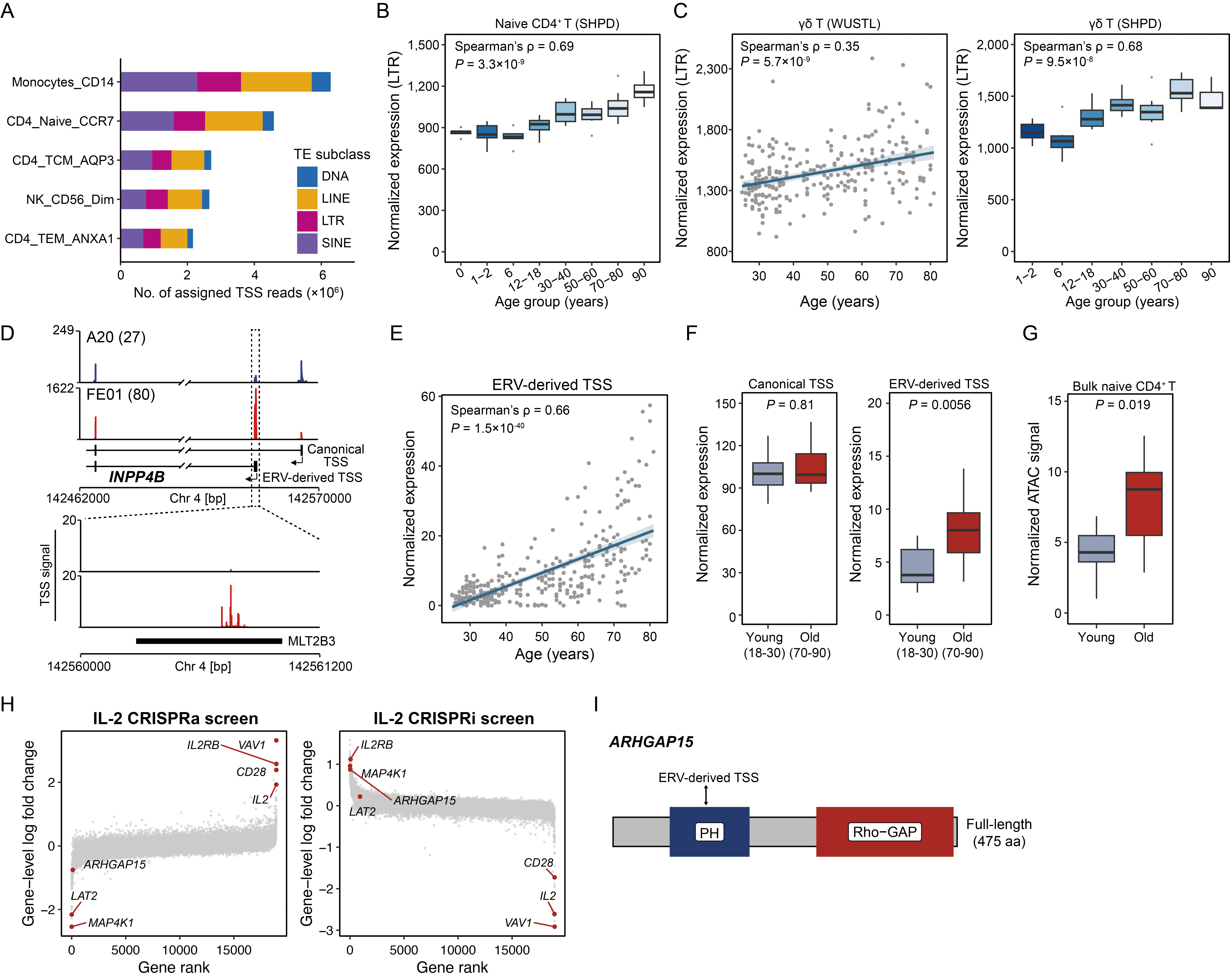

**Supplementary figure 8. Age-associated transcriptional activation of transposable elements.** (A) Bar plot showing the number of uniquely mapped TSS reads assigned to major transposable element classes across five immune cell types. (B) Normalized expression of LTR elements with age in CD4+ naïve T cells (SHPD cohort). (C) Age-associated upregulation of aggregated LTR expression in γδ T cells, validated across both WUSTL (left) and SHPD (right) cohorts. (D) Top: Pseudobulk scRNA-seq tracks of CD4+ naïve T cells illustrating elevated expression of a truncated *INPP4B* isoform drived by an ERV-derived promoter in old individuals. Bottom: Normalized TSS signals supporting the activation of ERV-derived TSS. (E) Correlation between age and the expression of the ERV-derived short *INPP4B* isoform in CD4+ naïve T cells (WUSTL cohort). (F) Comparison of normalized expression levels for the canonical long isoform (left) and the ERV-derived short isoform (right) of *ARHGAP15* in CD4+ naïve T cells between young and old individuals (SHPD cohort). (G) Increased chromatin accessibility at the ERV-derived promoter of the short *ARHGAP15* isoform, based on bulk ATAC-seq data of isolated CD4+ naïve T cells from different individuals. (H) Genome-wide CRISPR screens identify regulators of IL-2 production in primary human CD4+ T cells. Scatter plots depict the median sgRNA log2-fold change between high and low IL-2-producing cell populations. Results from the CRISPRa (activation) screen are shown on the left, and results from the CRISPRi (inhibition) screen are shown on the right. (I) Protein domain architecture of the *ARHGAP15* full-length isoform. Arrow indicates the position of the ERV-derived alternative TSS, which initiates the N-terminally truncated transcripts.

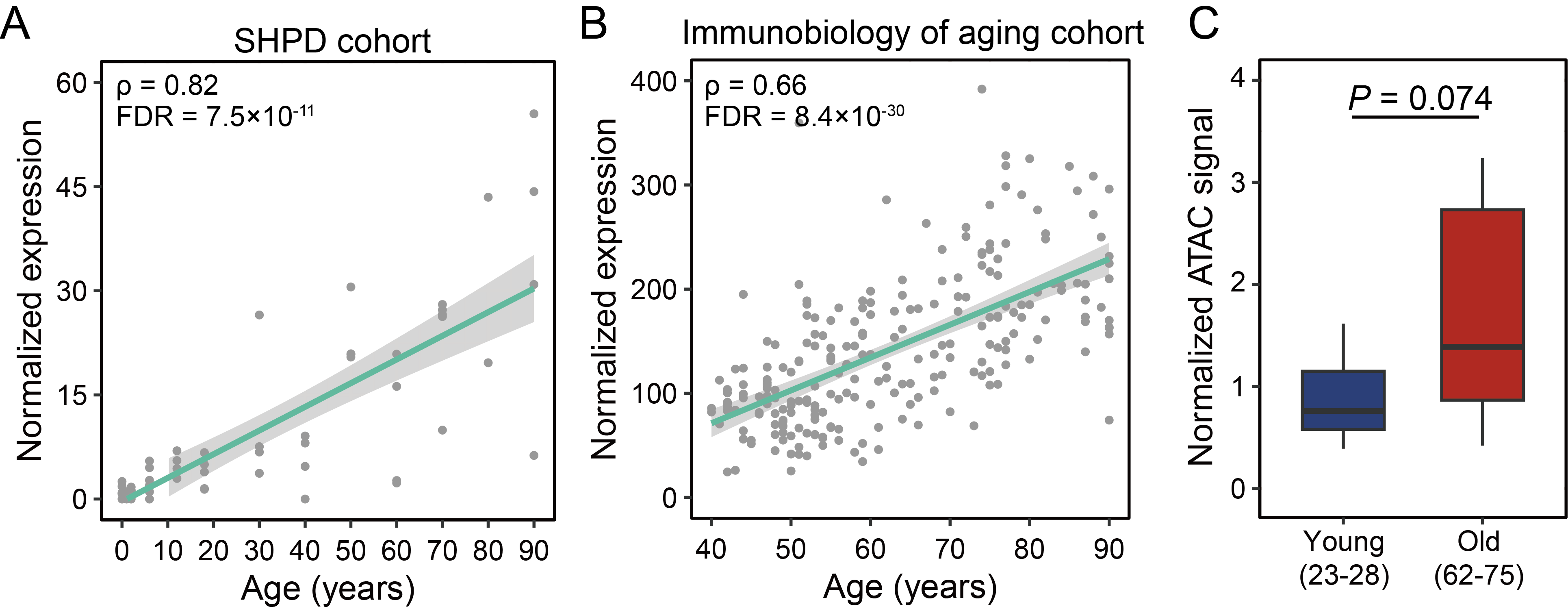

**Supplementary figure 9. Age-associated dynamics of *EDA* expression in CD4+ naïve T cells.** (A) Age-associated upregulation of *EDA* in CD4+ naive T cells in the SHPD cohort. (B) Age-associated upregulation of *EDA* in CD4+ naive T cells in a cohort of 234 healthy adults ranging in age from 40 to 90+ years old (Immunobiology of aging cohort). (C) Increased chromatin accessibility at the promoter region of *EDA*, based on bulk ATAC-seq data of isolated CD4+ naïve T cells from donors with different age.

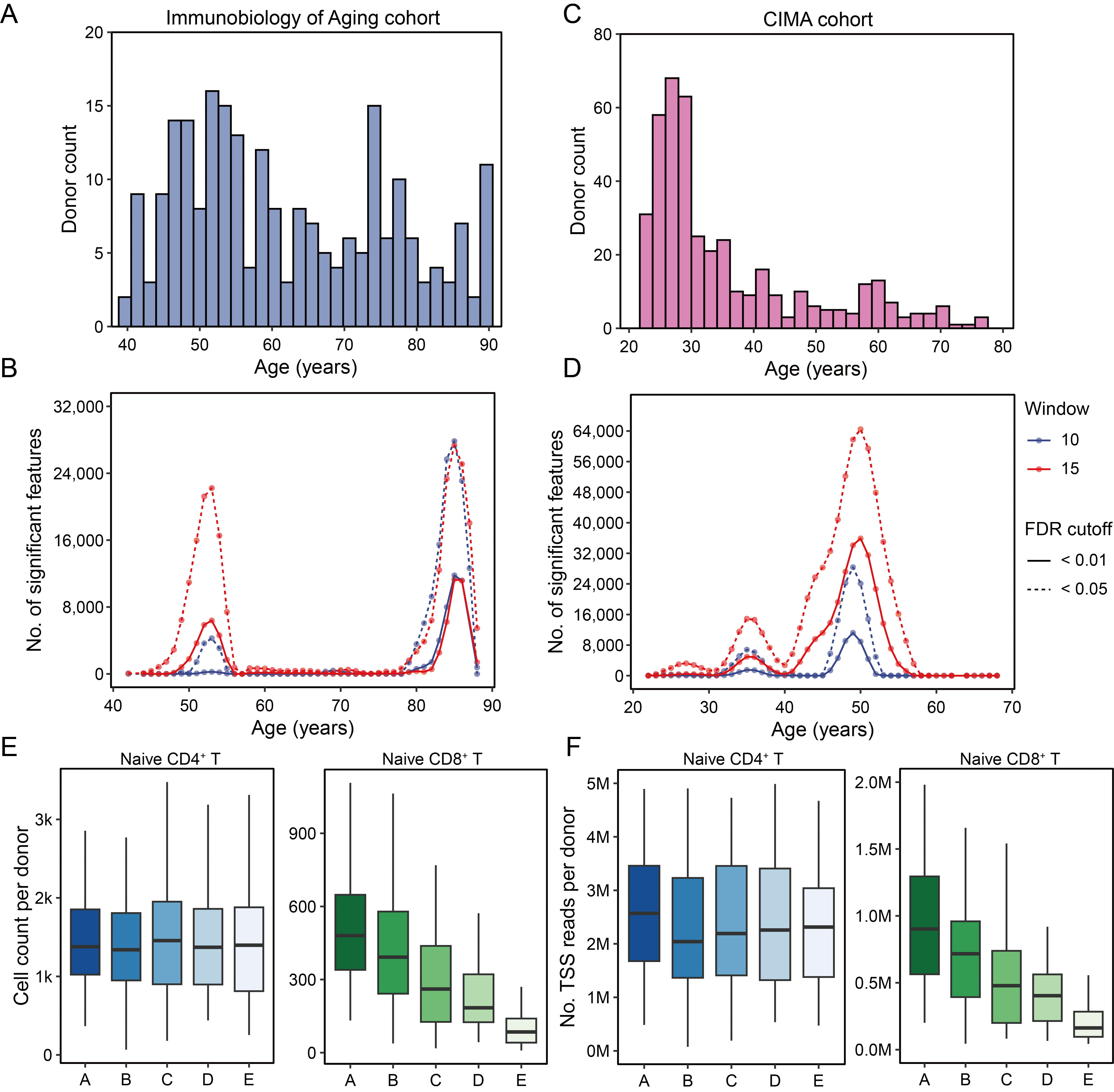

**Supplementary figure 10. Nonlinear dynamics of gene expression in immune aging and cohort data presentation.** (A, C) Distribution of chronological age across donor in the Immunobiology of Aging cohort (A) and the CIMA cohort (C). (B, D) Number of genes exhibiting significant age-associated expression changes across all immune cell types, as identified by DE-SWAN analysis in the Immunobiology of Aging cohort (B) and the CIMA cohort (D). For each cohort, the similar waves of aging were detected using different FDR thresholds and window size cutoffs. (E, F) Boxplots showing the number of sequenced cells (E) and the total number of TSS reads (F) per donor, stratified by cell type (naïve CD4⁺ T cells and naïve CD8⁺ T cells) and age group. Donors in the WUSTL cohort are categorized into five age groups: A (25-35), B (35-45), C (45-55), D (55-65), and E (65-81 years).
